# Potentiation of active locomotor state by spinal-projecting serotonergic neurons

**DOI:** 10.1101/2024.09.26.615260

**Authors:** Sara J. Fenstermacher, Ann Vonasek, Anne Cavanagh, Hannah Gattuso, Corryn Chaimowitz, Thomas M. Jessell, Susan M. Dymecki, Jeremy S. Dasen

## Abstract

Animals produce diverse motor actions that enable expression of context-appropriate behaviors. Neuromodulators facilitate behavioral flexibility by altering the temporal dynamics and output of neural circuits. Discrete populations of serotonergic (5-HT) neurons target circuits in the brainstem and spinal cord, but their role in the control of motor behavior is unclear. Here we define the pre- and post-synaptic organization of the spinal-projecting serotonergic system and define a role in locomotor control. We show that while forebrain-targeting 5-HT neurons decrease their activity during locomotion, subpopulations of spinal projecting neurons increase their activity in a context-dependent manner. Optogenetic activation of ventrally projecting 5-HT neurons does not trigger initiation of movement, but rather enhances the speed and duration of ongoing locomotion. We find that serotonergic neurons can influence motor output beyond periods of increased activity, indicating neuromodulators can act in the motor system over extended time scales. These findings indicate that the descending serotonergic system potentiates locomotor output and demonstrate a role for serotonergic neurons in modulating the temporal dynamics of motor circuits.

## Introduction

Neuromodulators act throughout the central nervous system to enable behavioral flexibility in changing environments^1,2^. Neuromodulators alter the dynamics of neural circuits, often leading to profound changes in network output. In the motor system, rhythm-generating circuits that produce essential behaviors such as walking, breathing, and swimming, are under strong neuromodulatory control^3,4^. While studies of rhythm-generating circuits have yielded insights into the mechanisms of neuromodulation, how neuromodulators are integrated with instructive motor commands and effector neurons to control behavior is less understood.

The monoamine serotonin (5-HT) is an evolutionarily-conserved modulator of motor circuits, including the rhythmically active circuits required to produce locomotion^5–21^. In vitro and pharmacological manipulations have shown serotonin is a potent modulator of spinal motor circuits^22^. Application of 5-HT to ex vivo neonatal spinal cord induces rhythmic activity similar to that observed during locomotion^14,23^. In vivo, 5-HT agonists and antagonists significantly alter the magnitude and timing of muscle activity during locomotion, as well as the strength of spinal reflexes^7,15,24,25^. At the cellular level, 5-HT promotes the excitability of spinal motor neurons (MNs) and ventral interneurons (INs)^26,27^, in part mediated by the generation of calcium dependent plateau potentials^11,18,28^. Despite a rich history of investigating 5-HT actions in the spinal cord, a major challenge has been to resolve how descending serotonergic pathways operate during movement and to determine their role in locomotor behavior.

The majority of serotonergic neurons reside within the raphe nuclei near the midline^29,30^ and are divided into two discrete clusters: a rostral group which largely projects to the forebrain and a caudal group that projects to the brainstem and spinal cord^31^. Within the caudal group, raphe obscurus (ROb) and raphe pallidus (RPa) project into ventral regions of the spinal cord, while raphe magnus (RMg) targets dorsal laminae^32–38^. Recent studies combining genetic fate mapping and molecular profiling have defined multiple anatomically and functionally distinct serotonergic neuron populations^37–41^. In particular, genetically-distinct serotonergic nuclei in the ventral medulla target either sensory or motor regions with the brainstem and spinal cord. Serotonergic neurons with a history of expressing the transcription factor encoding gene *Egr2* project dorsally while those expressing the neuropeptide-encoding gene *Tac1* terminate ventrally^37,38^.

Physiological studies revealed that the activity of 5-HT neurons correlates with movement. Extracellular recordings in cat showed that the firing rates of ROb and RPa neurons increase during treadmill-induced locomotion, indicating the activity of spinal-projecting 5-HT neurons is highly correlated to motor activity. By contrast, these neurons are relatively silent during REM sleep, a behavioral state characterized by cessation of movement and decreased muscle tone^42^. These observations suggest that descending modulation of spinal circuits by serotonergic neurons is involved in regulating motor function. Whether spinal-projecting serotonergic neurons affect the initiation, intensity, or duration of locomotor output is unresolved.

Here, we define the cellular and synaptic architecture of spinal cord-projecting serotonergic neurons and examine their function in motor control. We find that ROb/Pa provides direct input to spinal MNs and receives input from locomotor control regions, indicating the descending serotonergic system is recruited in parallel with motor command systems. The activity of ROb/Pa neurons correlates directly with locomotor activity and is distinct in nature and timing from other serotonergic populations. We show that activation of ROb/Pa neurons targeting ventral spinal circuits produce long-lasting increases in running duration and speed. Our studies reveal that anatomically and functionally distinct serotonergic neuron populations targeting ventral spinal motor circuits act to regulate locomotor behavior over long time scales.

## Results

### Spinal motor circuits receive biased 5-HT input from medullary raphe nuclei

To examine the spinal cord targets of subtypes of medullary serotonergic neurons, we first analyzed the distribution of spinal cord 5-HT in adult mice. Punctate 5-HT immunoreactivity was detected throughout the rostrocaudal extent of the spinal cord, with strongest labeling observed in the superficial dorsal horn, intermediolateral column, and ventral horn (Figures 1A, 1B, and 1C), confirming earlier work^29^. In the ventral horn, serotonin fibers densely targeted ChAT^+^ motor neurons, including both limb-innervating lateral motor column (LMC) (Figure 1D) and axial-innervating medial motor column (MMC) neurons (Figure S1B). Serotonin puncta were also observed in proximity to the cell bodies of ventral interneurons, including excitatory Chx10^+^ V2a neurons (Figure 1E), and V0c interneurons near the central canal (Figure S1A).

**Figure 1.**
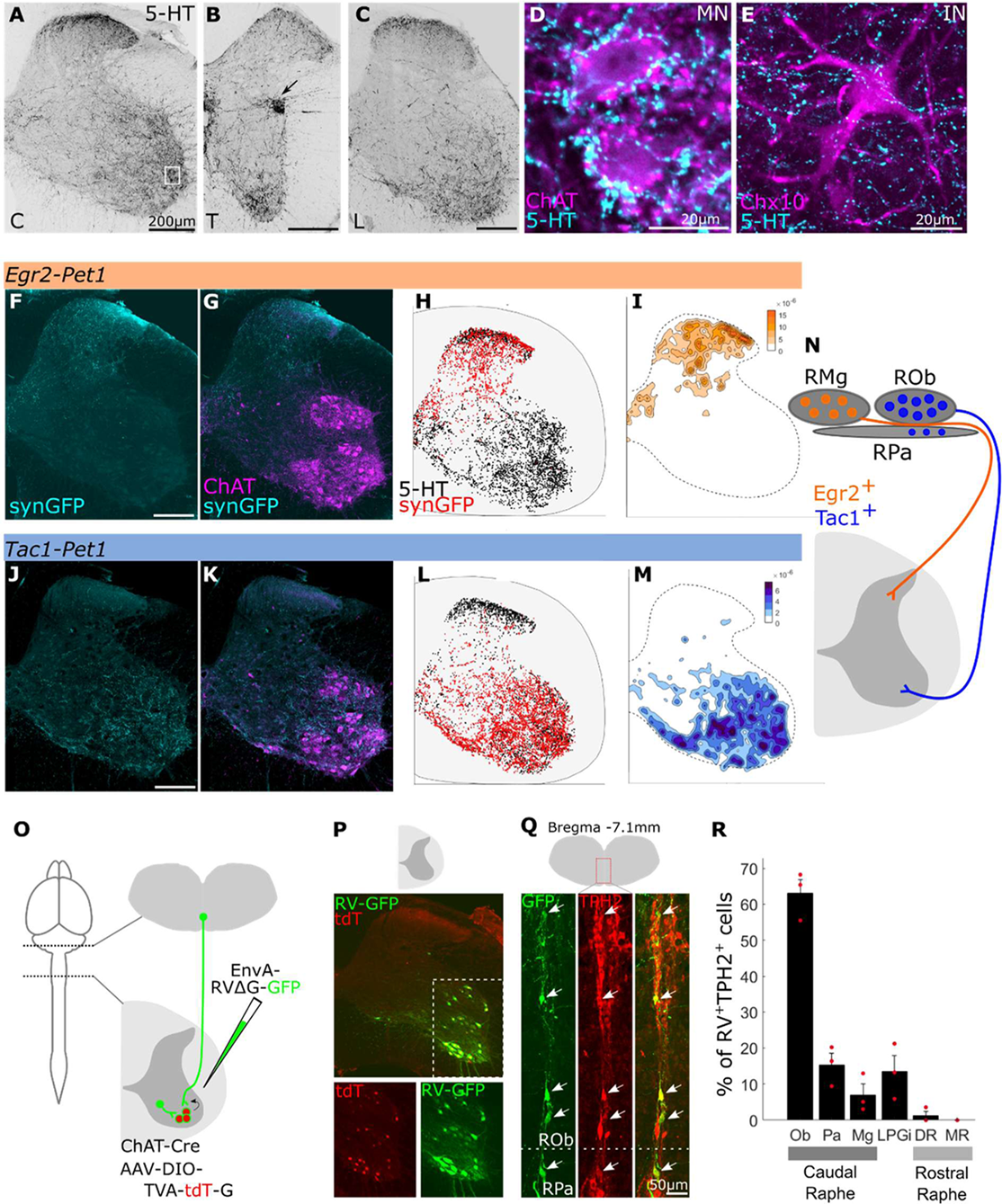
Spinal motor circuits receive biased 5-HT input form medullary raphe nuclei (A-E) 5-HT immunostaining in adult spinal cord at cervical (C), thoracic (T), lumbar (L) levels. Arrow in (B) shows dense 5-HT immunoreactive region at thoracic levels is around preganglionic neurons **(D)** ChAT^+^ motor neurons (MN) with 5-HT immunostaining. Zoom-in of white box in A. Scale bar 20µm. **(E)** Genetically labeled Chx10^+^ ventral interneuron (IN) with 5-HT immunostaining. Scale bar 20µm. **(F-I)** Genetic labeling of *Egr2-Pet1* neurons with synaptophysinGFP (cyan) in cervical spinal cord. **(G)** Immunostaining for ChAT (magenta) to visualize MNs. **(H)** Distribution of *Egr2-Pet1* puncta (red) and total 5-HT puncta (black). **(I)** Relative density plot of *Egr2-Pet1* puncta shown in F. **(J-M)** Genetic labeling of *Tac1-Pet1* neurons with synaptophysinGFP in cervical spinal cord. **(K)** Immunostaining for ChAT to visualize MNs. **(L)** Distribution of *Tac1-Pet1* puncta (red) and total 5-HT puncta (black). **(M)** Relative density plot of *Tac1-Pet1* puncta shown in J. Data in panels F-M are representative of at least 3 animals for each genotype. **(N)** Schematic of *Tac1* vs. *Egr2 Pet1* neuron populations targeting spinal cord. **(O-R)** Monosynaptic retrograde rabies tracing to identify 5-HT input to spinal MNs. **(O)** Experimental procedure. EnvA-RVΔG-GFP injection into spinal cord (C4-T1) to infect spinal MN expressing TVA and G protein. **(P)** Cervical spinal cord (C8) MNs infected with AAV-DIO-TVA-tdT-G and rabies virus (RV). Zoom-ins of white-dotted region are below. **(Q)** Retrogradely labeled neurons expressing tryptophan hydroxylase 2 (TPH2) within ROb and RPa. Arrows indicating RV^+^/TPH2^+^ cells. Scale bar 50µm. **(R)** Quantification of RV-labeled TPH2^+^ neurons. Percentage of total RV^+^/TPH2^+^ cells within raphe and LPGi. n=3, each red dot is average of a single animal. Error bars are SEM.

Building from prior serotonergic neuron subtype identification and efferent mapping^37,38^, we performed anterograde labeling of genetically defined 5-HT populations to visualize serotonergic projections to the spinal cord. We used an intersectional genetic-approach to visualize synaptophysinGFP-labelled terminals^43^ of two caudal 5-HT populations defined by co-expression of *Pet1* with *Egr2*^37^ or *Tac1*^38^ (Figures S1C and S1D). We observed that for *Egr2-Pet1* neurons, whose soma reside predominantly in Raphe Magnus (RMg)^37^ (Figure S1F), synaptophysinGFP-labeled terminals located in the dorsal horn of cervical, thoracic, and lumbar segments (Figures 1F, 1G, 1H, 1I, and S1E). By contrast, *Tac1-Pet1* neurons, whose soma reside within the Raphe Obscurus (ROb), Raphe Pallidus (RPa), and Lateral Paragigantocellularis (LPGi) nuclei of the caudal medulla^38^ (Figure S1F), selectively target the ventral horn throughout rostrocaudal levels and densely surround MNs (Figures 1J, 1K, 1L, 1M, and S1E). Genetically-distinct sets of *Pet1* neurons, therefore innervate non-overlapping domains along the dorsoventral axis of the spinal cord (Figure 1N), likely providing separate channels for serotonergic modulation of sensory input and motor output.

While neuromodulators such as serotonin are often released extrasynaptically to signal via volume transmission, MNs receive numerous 5-HT synaptic contacts upon their soma and dendrites^44^. To determine the origin of this direct and synaptic serotonergic neuron input to MNs, we performed retrograde tracing from spinal MNs using G-deleted rabies virus^45^, which travels exclusively through synapses (Figure 1O). To restrict expression of the avian TVA receptor and rabies glycoprotein to spinal MNs we injected AAV-DIO-TVA-tdT-G into the lateral ventricle of *Chat-Cre* mice at P0 (Figure 1P and S2A)^46^. Eight weeks later, we infected spinal MNs by delivery of EnvA pseudotyped, G-deleted N2c rabies virus with GFP (EnvA-N2cΔG-GFP) into the spinal cord (Figures 1O and 1P). We identified serotonin neurons by presence of tryptophan hydroxylase 2 (TPH2) and quantified GFP^+^TPH2^+^ neurons in the brainstem (Figures 1Q, S2B, and S2C). Rabies-labeled TPH2^+^ cells were found almost exclusively within the medullary raphe (ROb, RPa, RMg) and lateral paragigantocellularis (LPGi) nuclei, with over 60% of GFP^+^TPH2^+^ cells located in ROb (Figure 1R and S2D). Over 75% of GFP^+^TPH2^+^ cells were found within the neighboring ROb and RPa nuclei, thus we will henceforth refer to this population as ‘ROb/Pa’. Only a few cells were labeled in the dorsal raphe nucleus (DRN, Figure 1R). By comparison, similar transsynaptic tracing assays from V2a INs (using *Chx10-Cre* mice), labeled very few 5-HT neurons (Figures S2E-S2J). These results suggest that excitatory spinal INs do not receive substantial direct synaptic input from serotonergic neurons, despite local innervation, although we cannot rule out the possibility that there are differences in efficiency of RV transmission between INs and MNs.

We next examined whether serotonergic neurons target specific spinal populations, or instead broadly innervate multiple segments. We injected AAV2r-FLEX-tdTomato into the cervical spinal cord of *Pet1-Cre* mice to retrogradely label serotonergic neurons (Figure S3A). Following this, we observed tdTomato^+^ fibers throughout both thoracic and lumbar levels, demonstrating that descending 5-HT neurons exhibit a highly collateralized structure with widespread targets across rostrocaudal spinal segments (Figures S3B-S3D). Within lumbar segments we observed increased 5-HT density at specific ventral MN pools (Fig S3E and S3F). Together, these studies showed that within the spinal cord, fibers from ROb and Pa serotonergic neurons target neurons in the ventral horn, provide direct input to spinal MNs, and that these inputs distribute to more than one axial level.

### Activity of ventrally-projecting serotonin neurons increases during locomotion

The activity of serotonergic neurons can depend on behavioral context, with specific populations increasing or decreasing their activity during movement^42,47,48^. However, the dynamics of specific serotonergic populations during locomotion are just beginning to be defined. We therefore measured the activity of three serotonergic neuron populations (ROb/Pa, RMg, or DRN) in mice during both wheel and treadmill running. To record the activity of specific 5-HT subpopulations, we performed fiber photometry in *Pet1-Cre* transgenic mice that were infected to express the calcium indicator GCaMP6s (AAV-FLEX-GCaMPs6, Figures 2C, 2G, 2K, S5B, S5D, and S5F). One week following viral injection, a low-profile running wheel was placed into each mouse cage for introduction and practice during the remainder of the experiment. One week later, mice were acclimated to the recording area and allowed to run on the wheel and treadmill. Starting 3 weeks after the viral injection, photometry signals were recorded during 30-minute sessions on the wheel (Figures 2A and 2B) and for 10s intervals on the treadmill with delivery of 470nm excitation light to monitor neural activity and 405nm control light to assess motion artifact.

**Figure 2.**
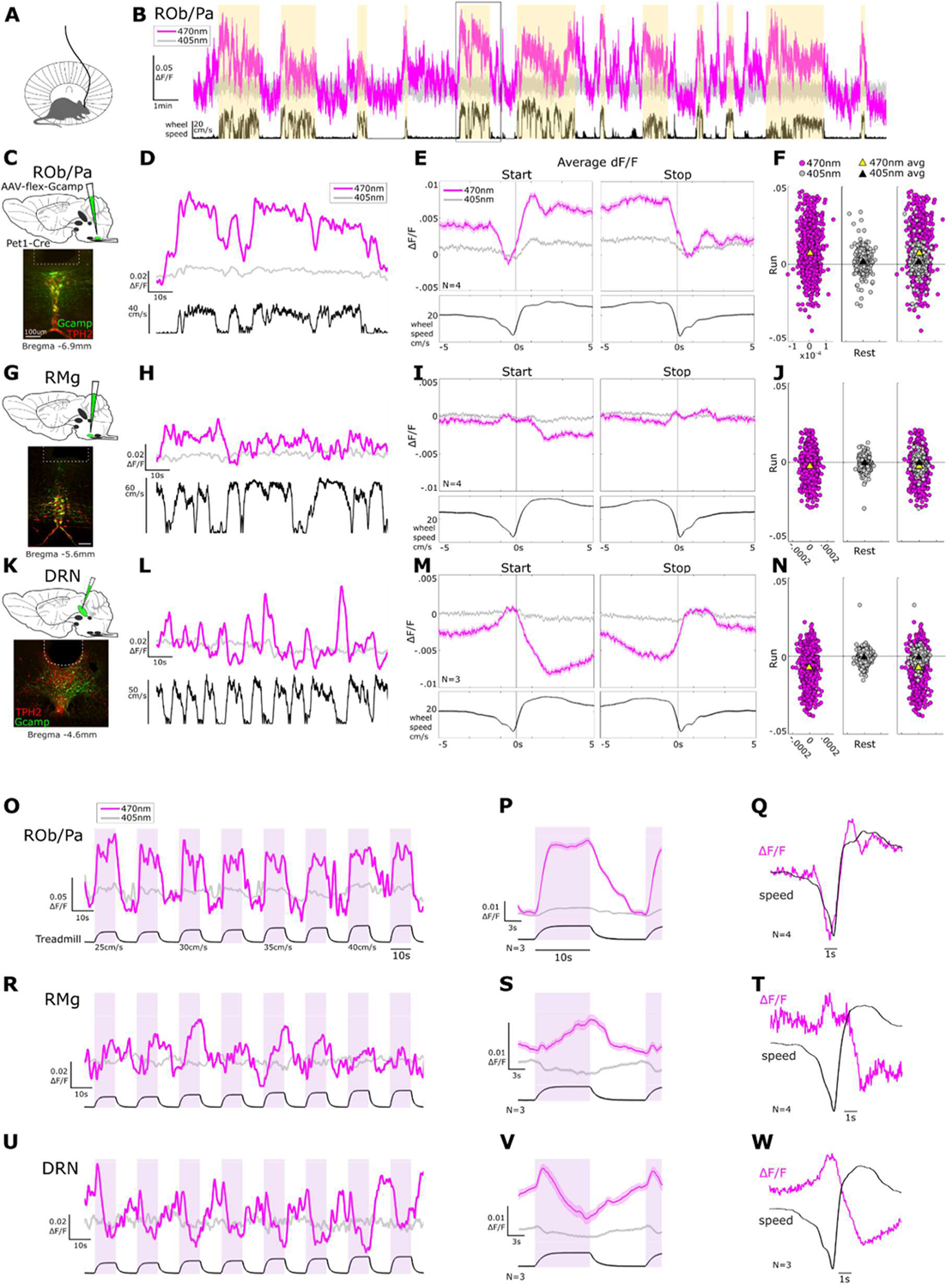
Activity of ventrally-projecting serotonin neurons increases during locomotion. **(A)** Behavioral assay. Mice freely running on low-profile running wheel with attached fiber for photometry recordings. **(B)** Example photometry traces ΔF/F (470nm in pink, 405nm in grey) from ROb/Pa *Pet1* neurons with wheel speed (black) during single session. **(C,G,K)** Injection of AAV-FLEX-Gcamp6s into *Pet1-Cre* mice to infect *Pet1* neurons. Histology showing Gcamp6s expression and TPH2 immunostaining in ROb/Pa **(C)**, RMg **(G)**, and DRN **(K)**. **(D,H,L)** Example trace during 60sec of free running with ΔF/F and wheel speed for ROb/Pa *Pet1* neurons (**D**), RMg *Pet1* neurons (**H**), DRN *Pet1* neurons (**L**). **(E,I,M)** Averaged ΔF/F and wheel speed aligned to start and stop of running. ROb/Pa 1252 runs (n=4 mice), RMg 1290 runs (n=4 mice), DRN 1172 runs (n=3 mice). **(F,J,N)** Plots of the average ΔF/F during the rest (prior to run start) on the x-axis and during the run on the y-axis at 470nm (pink) and 405nm (grey). Each dot is an individual run. Triangles show average ΔF/F of all runs at 470nm (yellow) and 405nm (black). Average run ΔF/F for ROb/Pa 0.0072, 0.0016 (470nm,405nm); RMg -0.0025, -0.0005; DRN - 0.0073, -0.0008. For ROb/Pa we do not observe a correlation between neural activity during wheel running with running speed (Figure S5G). **(O,R,U)** Example photometry traces during single run of treadmill assay showing 470nm (pink) and 405nm (grey) ΔF/F with treadmill speed in black. ROb/Pa *Pet1* neurons (**O**), RMg *Pet1* neurons (**R**), DRN *Pet1* neurons (**U**). **(P,S,V)** Averaged signal across all treadmill run bouts with 10sec run and 10sec rest. ROb/Pa 120 trials, n=3 mice (**P**), RMg 120 trials, n=3 mice (**S**), DRN 120 trials, n=3 mice (**V**). **(Q,T,W)** Average 470nm ΔF/F (pink) and wheel speed (black) at run start on wheel. ROb/Pa (**Q**), RMg (**T**), DRN (**W**).

We observed that ROb/Pa *Pet1* neurons display a sustained increase in activity during periods of running (Figures 2B, 2D, and S4A). Averaged ΔF/F during the start and stop of run bouts shows an increase in activity at running onset and decrease at offset (Figures 2E, 2Q, S5A and S5H). A scatterplot of averaged ΔF/F values during run bouts vs. rest periods just prior to the run shows the neural activity increased during locomotion (Figure 2F). Similarly, we observed a robust increase in GCaMP signal during treadmill-evoked locomotion (Figures 2O and 2P), as was observed by Veasey et al.^42^. During both spontaneous and evoked locomotion, ROb/Pa *Pet1* neurons displayed a rapid increase activity at the onset of movement (Figures 2P and 2Q). Thus, ROb/Pa *Pet1* neurons that project to the spinal ventral column exhibit activity that positively correlates with locomotion.

By contrast, the activity of DRN *Pet1* neurons were anti-correlated with locomotion. DRN activity decreased during running (Figure 2L and S4C), with a consistent reduction in signal at the start of running and an increase in activity when the animal stops (Figures 2M, 2W, and S5E). The scatterplot of averaged ΔF/F values during run bouts vs. rests shows the neural activity during run bouts shifted below zero, reflecting decreased activity during locomotor bouts (Figure 2N). We also observed a similar decrease in DRN activity during treadmill locomotion (Figures 2U and 2V). Thus, DRN and ROb/Pa *Pet1* neurons display reciprocal patterns of activity during locomotion.

Finally, to determine whether movement-correlated activity in ROb/Pa is unique to ventral spinal cord-projecting 5-HT neurons, we compared GCaMP signal to the neighboring dorsal spinal cord-projecting RMg (Figure 2G). We observed that RMg *Pet1* neuron activity did not strongly correlate with locomotion when animals ran freely on the wheel (Figures 2H,2I, 2J, 2T, S4B, and S5C). When animals were forced to perform treadmill running however, RMg *Pet1* neuron activity was positively correlated to movement, with a slow rise in activity during locomotion and decrease during rest (Figures 2R and 2S), suggesting the activity of RMg neurons during movement depends upon context. Altogether, these observations indicate that the activity of serotonergic neurons that send their projections ventral spinal motor circuits is distinct from 5-HT populations targeting dorsal spinal cord and brain regions.

### ROb/Pa activity is a strong predictor of locomotor state

To determine whether the activity of serotonin neurons could be used to predict behavioral state, we used a linear-non-linear model to examine the relationship between neural activity and locomotor behavior^49,50^ We generated linear filters for ROb/Pa, RMg, and DRN, which illustrate the temporal relationship between neural activity and wheel speed (Figure 3). This analysis was performed using total wheel running data to assess whether the patterns of activity observed during running onset and offset extend throughout running behavior. Indeed, the model generated unique filters for each of the three serotonergic populations that were consistent with the activity patterns we observed in the previous analysis (Figure 2).

**Figure 3.**
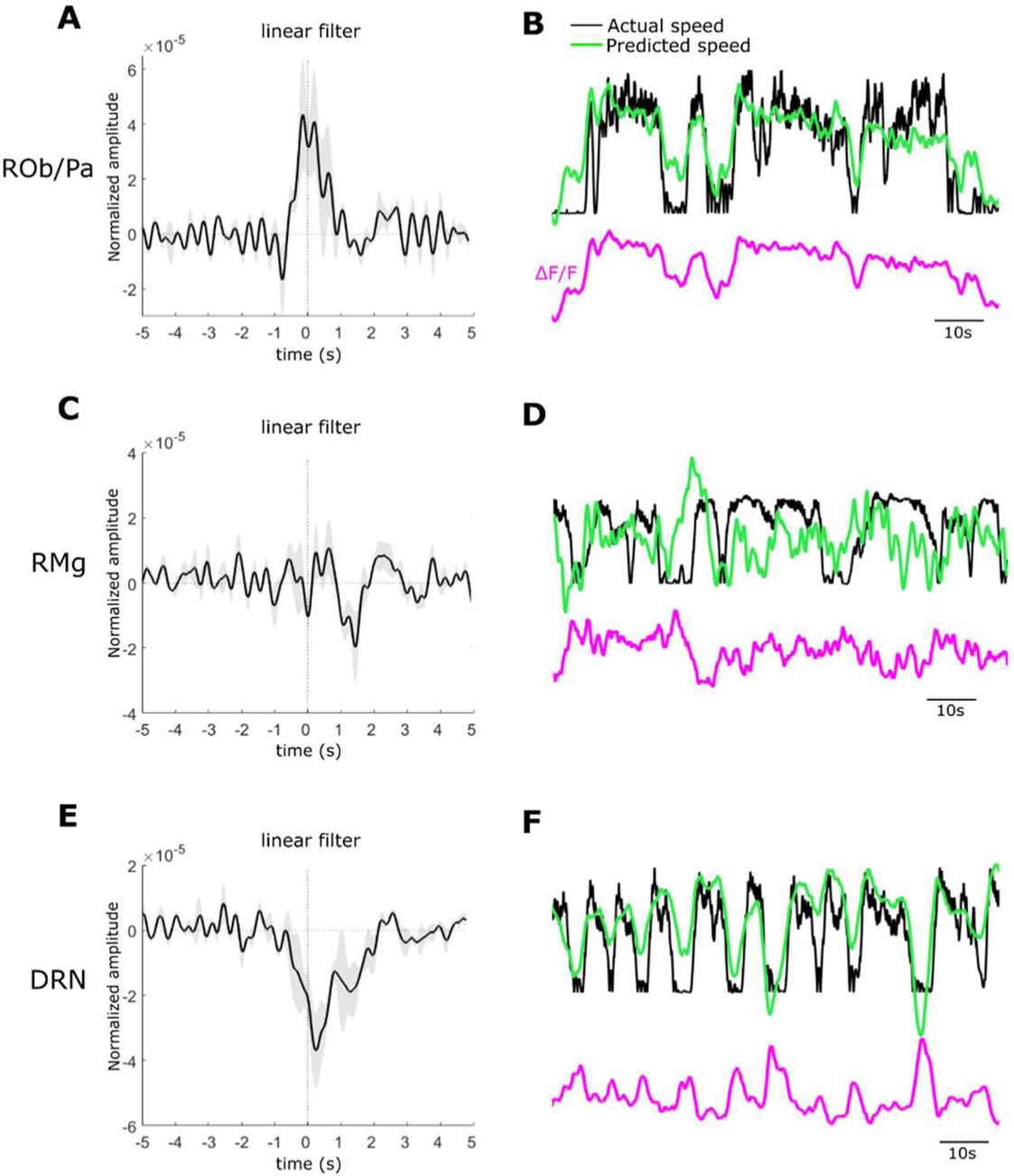
Locomotor state is a strong predictor of ROb/Pa activity (A,C,E) Linear filters for ROb/Pa *Pet1* neurons (**A**), RMg *Pet1* neurons (**C**), DRN *Pet1* neurons (**E**). Black line is average (n=3 mice) with grey SEM. **(B,D,F)** Representative traces for actual wheel speed (black) overlayed with predicted speed (green) and aligned ΔF/F trace below (pink) for ROb/Pa (**B**), RMg (**D**), DRN (**F**).

The ROb/Pa filter contains a single, positive peak, indicating that increases in wheel speed occur during increases ROb/Pa *Pet1* neuron activity (Figure 3A). By contrast, the DRN filter contains a negative peak (Figure 3E), indicating that increases in wheel speed occur during decreases DRN *Pet1* neuron activity. The position of each peak around 0 implies that changes in ROb/Pa and DRN *Pet1* neuron activity are coincident to changes in speed, as we observed in the aligned ΔF/F and wheel speed traces (Figures 2Q and 2W). To assess the quality of these models, we used each filter to predict locomotor speed from neural activity and compared this to the actual wheel speed (Figures 3B and 3F) We find that ROb/Pa *Pet1* neuron activity (Figure 3B, CC=0.35, SEM=0.11) and DRN *Pet1* neuron activity (Figure 3F, CC=0.29, SEM=0.02) are good predictors of wheel running.

By comparison, the RMg filter is noisier and contains less pronounced structure (Figure 3C), likely due to the varied signal we observed during wheel running (Figure 2H). We find using this model that RMg *Pet1* neuron activity is a poor predictor of wheel speed (Figure 3D, CC=0.14, SEM=0.06). The RMg filter does however contain a small negative peak right of 0, suggesting that changes in speed precede change in RMg *Pet1* neuron activity. This is consistent with the slower RMg dynamics we observed during both wheel and treadmill locomotion (Figures 2S and 2T). Thus, in comparison to other 5-HT populations, ROb/Pa exhibits a distinct pattern of activity that is highly correlated to locomotor activity.

### Brain-wide inputs to raphe *Pet1* neurons targeting ventral spinal cord

The tight coupling between ROb/Pa *Pet1* neuron activity and locomotion suggests that ROb/Pa neurons may be part of a broader network involved in locomotor control. Descending command pathways within the midbrain and brainstem are known to regulate the start, stop and speed of locomotion^51–54^. The cuneiform nucleus (CnF), a component of the mesencephalic locomotor region (MLR), is important for regulating locomotor initiation and speed^52,53^. The brainstem LPGi, acts directly upon spinal locomotor networks to control motor output^51^. To determine whether ROb/Pa neuron neurons receive input from these locomotor control areas, we performed monosynaptic rabies tracing to identify brain regions that provide direct input to ventrally-projecting serotonin neurons (Figures 4A and 4B).

**Figure 4.**
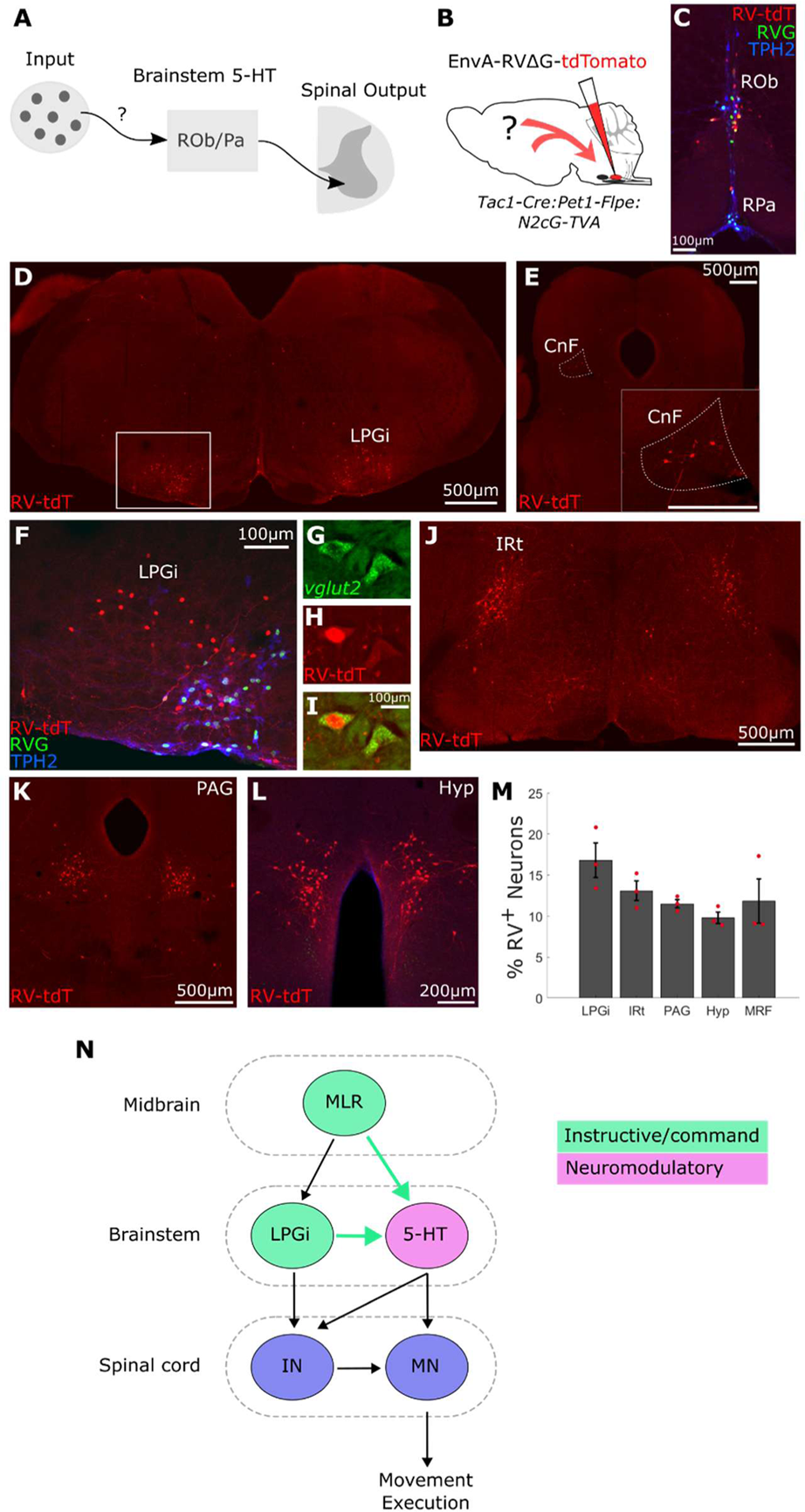
Brain-wide inputs to raphe *Pet1* neurons targeting ventral spinal cord. **(A)** Schematic showing input pathways to 5-HT neurons targeting the ventral spinal cord. **(B)** Experimental procedure. Injection of EnvA-RVΔG-tdTomato into ROb/Pa of *Tac1-Cre:Pet1-Flpe:N2cG-TVA* mice to identify monosynaptic inputs to *Tac1-Pet1* neurons. **(C)** Primary infection of *Tac1-Pet1* neurons in ROb/Pa with EnvA-RVΔG-tdTomato. **(D-L)** Representative images of EnvA-RVΔG-tdTomato (tdT) infected neurons LPGi (**D**, zoom-in white box in **F**), cuneiform (CnF, **E**), intermediate reticular nucleus (IRt, **J**), periaqueductal grey (PAG, **K**), hypothalamus (Hyp, **L**). **(G-I)** *Vglut2* mRNA expression with EnvA-RVΔG-tdTomato infection in LPGi neurons. Panel I shows overlay of *Vglut2* mRNA and RV-tdT. **(M)** Percentage of second-order rabies-infected neurons in various brain areas out of total rabies-labeled cells (n=3), each red dot is percentage from single animal. Error bars are SEM. **(N)** Model suggested by rabies tracing. Locomotor command neurons within MLR (CnF) and LPGi send projections (cyan arrows) to brainstem 5-HT neurons that target ventral spinal cord to facilitate modulation of spinal MNs and INs during locomotor behavior. (lateral paragigantocellularis, LPGi; intermediate reticular nucleus, IRt; periaqueductal grey, PAG; hypothalamus, Hyp; medullary reticular formation, MRF; cuneiform, CnF; superior colliculus, SC; raphe pallidus, RPa; raphe obscurus, ROb; raphe magnus, RMg, mesencephalic locomotor region, MLR)

To restrict rabies infection to *Tac1-Pet1* ROb/Pa neurons, we used an intersectional strategy by generating a knock-in mouse line allowing for Cre- and Flpe-dependent expression of the TVA receptor and rabies N2c glycoprotein (Figures S6A and S6B). To label inputs to the *Tac1-Pet1* subpopulation, we injected *Tac1-Cre:Pet1-Flpe:N2cG-TVA* mice with N2c rabies virus (EnvA-N2cΔG-tdTomato) (Figure 4C). The rabies glycoprotein was tagged with HA, allowing for identification of the initially infected starter population (HA^+^tdT^+^)(Figure S6D). We observe no HA expression or RV infection in *N2cG-TVA* mice in the absence of Cre and Flpe (Figure S6C), indicating that rabies tracing depends on cell-type specific expression of N2cΔG and TVA.

Following rabies injection of *Tac1-Cre*:*Pet1-Flpe:N2cG-TVA* mice, we observed labeled tdT^+^ cells in several distinct brain areas. We identified dense bilateral clusters of tdT^+^ cells within LPGi, intermediate reticular nucleus (Irt), periaqueductal gray (PAG), and hypothalamus (Figures 4D, 4F, 4J, 4K, and 4L). Additionally, we observed sparse labeling within CnF (Figure 4E), the medullary reticular formation (MRF), including gigantocellularis reticular nucleus Gi, the ventral part (GiV), the alpha part (GiA), and the medullary reticular formation ventral part (MdV), superior colliculus, and sensorimotor cortex (Figure S6E and data not shown). We found comparable labeling of these regions after tracing from ROb/Pa neurons in *Pet1-Cre* mice (Figures S6F-S6I).

Several of the pre-ROb/Pa *Pet1* neuron populations, including CnF, LPGi, MRF and PAG, have known roles in motor control^51,52,55,56^. The highest percentage of rabies-labeled cells (∼16%) were located in LPGi (Figure 4M). Recent work identified Vglut2+ neurons within the LPGi that activate and are required for high-speed locomotion^51,57^. Therefore, we performed in situ hybridization for *vglut2* mRNA and identified Vglut2-expressing RV-labeled cells in LPGi (Figures 4G, 4H, and 4I). These observations suggest a model whereby descending serotonergic pathways are recruited in conjunction with locomotor command neurons to modulate spinal motor circuits (Figure 4N).

### Activation of ROb/Pa potentiates ongoing locomotor behavior

The activity of ROb/Pa *Pet1* neurons during wheel running and connectivity with multiple locomotor control regions suggests that serotonergic input may be important for regulation of locomotion. However, it is unknown whether spinal cord-projecting 5-HT neurons regulate specific features of locomotion, such as initiation, duration, speed, and termination. To test this, we measured the effect of optogenetically activating ROb/Pa neurons during wheel running. We injected Cre-dependent channel rhodopsin (AAV-DIO-ChR-EYFP) into ROb/Pa of *Pet1-Cre* mice (Figures 5A, S7A, and S7B). Two weeks post injection mice were acclimated to the wheel and fiber attachment. Three weeks post injection, mice were placed on running wheels for 30 minutes, during which they received alternating 5-minute periods of light delivery (5 sec 20Hz pulses) and no light (Figure 5B). Control Cre-negative littermates were injected with ChR virus and did not exhibit any ChR expression (Figure S7C). Animals spent about two thirds of their time on the wheel locomoting (Figure 5C).

**Figure 5.**
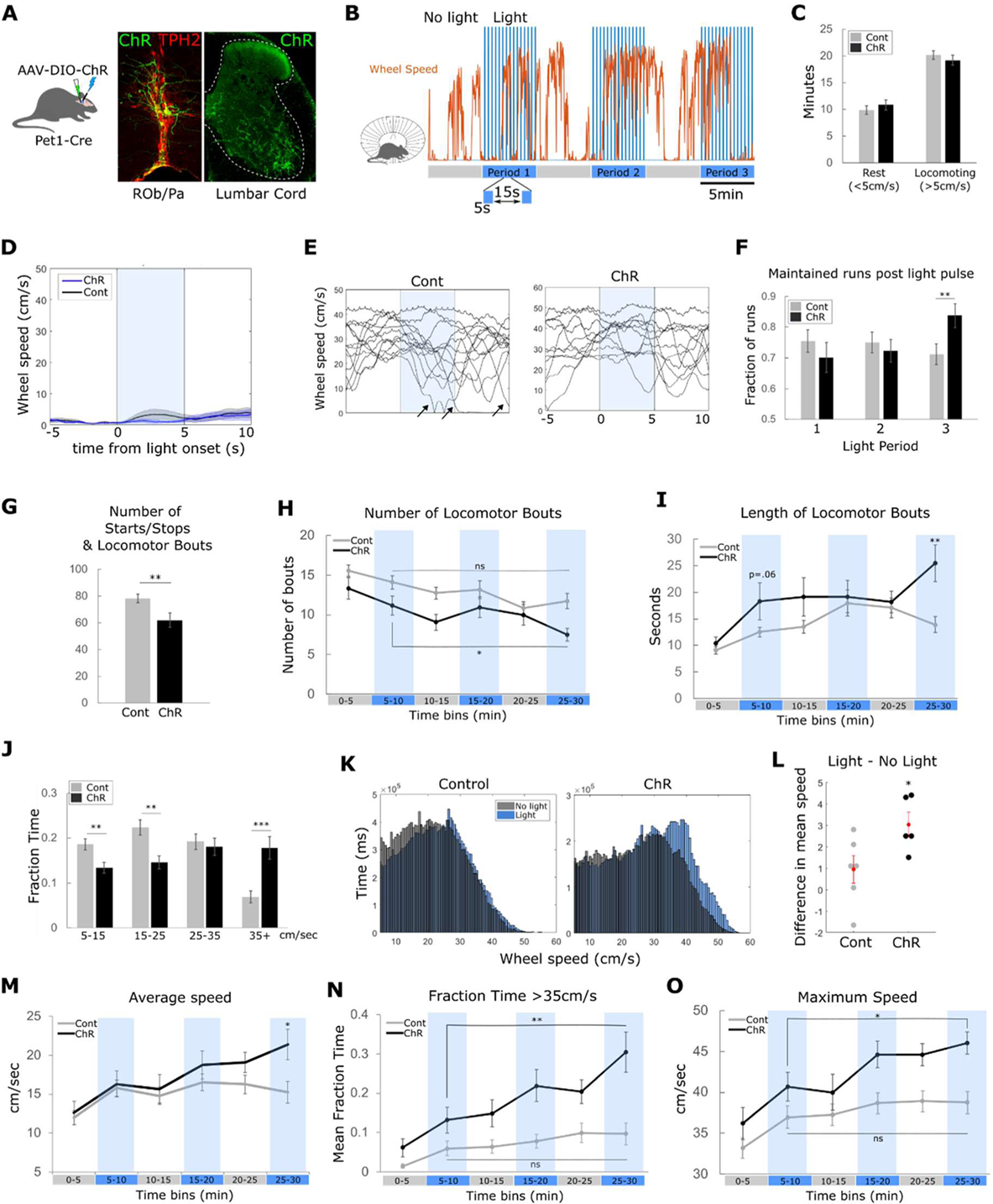
Activation of ROb/Pa potentiates ongoing locomotor behavior (A-B) Experimental procedure. **(A)** Injection of AAV2-DIO-ChR2(H134R)-EYFP into ROb/Pa of *Pet1-Cre* mice. ChR-expressing TPH2^+^ neurons in ROb/Pa with ChR^+^ axon terminals in lumbar spinal cord. **(B)** Light delivery protocol during 30min of wheel running with single example wheel speed trace from one ChR animal (orange). Stimulation during 5-minute light periods: 5s 20Hz pulses 470nm light repeating every 15s. **(C)** Total time at rest vs. locomoting in control (grey) and ChR (black) animals. n=22-30 trials from 5-6 animals. **(D)** Wheel speed for trials when animal was still at time of light onset. Averaged wheel speed from first light pulse of each light period for control and ChR animals when wheel was 0cm/s at time of light onset. **(E)** Example wheel traces from light period 3 when animal is running at time of light pulse. Arrows indicate trials when animals stop running. **(F)** Fraction of trials when animal is running at time of light onset and where running is maintained for 10 seconds following light onset. n=113-159 runs, from 5-6 animals. Error bars are SEM. *p<0.05 unpaired t-test. **(G)** Total number of locomotor starts, stops, and bouts for ChR and control mice n=22-30 trials from 5-6 animals. **p<0.01 unpaired t-test. **(H)** Number of locomotor bouts during each 5-minute light or no light period for control and ChR animals. n=22-30 trials from 5-6 animals. *p<0.05 unpaired t-test. ns=not significant. **(I)** Average length of locomotor bouts during each 5-minute light or no light period for control and ChR animals. n=22-30 trials from 5-6 animals. **p<0.01 unpaired t-test. **(J)** Fraction of total time control or ChR animals spend within various locomotor speed intervals. n=22-30 trials from 5-6 animals. **p<0.01, ***p<0.001 Bonferroni-corrected t-test. **(K)** Distribution of time (ms) across wheel speed during No-light (grey) or Light (blue) periods. Combined data from 5-6 animals across all No-light or Light periods. **(L)** Difference in average speed between Light and No-light periods. n=5-6 animals. *p<0.05 unpaired t-test. **(M)** Average speed during each 5-minute light or no light period for control and ChR animals. n=22-30 trials from 5-6 animals. *p<0.05 unpaired t-test. **(N)** Fraction of time mice spent locomoting greater than speed of 35cm/s during each 5-minute light or no light period for control and ChR animals. n=22-30 trials from 5-6 animals. **p<0.01 unpaired t-test. ns=not significant. **(O)** Average maximum speed during each 5-minute light or no light period for control and ChR animals. n=22-30 trials from 5-6 animals. *p<0.05 unpaired t-test. ns=not significant. All Error bars are SEM.

Because serotonin acts through both ionotropic and metabotropic receptors, changes in locomotor parameters might emerge over both short and long times scales. We therefore selected a pattern of blue light delivery that included both short 5-second pulses and longer 5-minute periods with or without light pulses to enable us to observe how neuronal activation influenced behavior over short and long-time scales. We did not observe locomotor initiation or acute changes in speed in any animals upon blue light stimulation of ROb/Pa *Pet1* neurons (Figures 5D, S7F, and S7G). This is in stark contrast to locomotor-driving regions CnF and LPGi, which when stimulated result in short latency activation of locomotion^51,52,57^. However, we observed that ChR animals exhibited an increase in the fraction of runs that were maintained following a light pulse (Figures 5E and 5F) and fewer total stops in locomotion (Figure 5G), suggesting that serotonergic input reduces the probability of terminating locomotor output.

We asked whether activation of ROb/Pa *Pet1* neurons influenced the amount of time animals spent engaged in locomotion (wheel speed >5cm/sec). On average, mice spent approximately 66% of their time moving on the wheel (∼20min), for both control and ChR animals (Figure 5C). Thus, activation of ROb/Pa *Pet1* neurons did not affect the total duration of locomotion. We next sought to determine whether activation of serotonergic neurons influenced the number of discrete locomotor bouts mice produced. Serotonin increases the excitability of spinal MNs^11,26^. Thus, we predicted that activation of ROb/Pa may lead to an increase in run bouts, as heightened MN excitability could lower the threshold needed for a command region to initiate locomotion. Unexpectedly, we observed the opposite outcome, as ChR animals produced fewer locomotor bouts than control animals (Figures 5G and 5H). However, since ChR animals run the same total amount of time as control animals (Figure 5C), this observation suggests that ChR animals run bouts are longer. Indeed, we found that activation of ROb/Pa *Pet1* neurons significantly increased the length of locomotor bouts by the third light period (Figure 5I). Together, these observations suggest that once an animal has initiated a run bout, ROb/Pa *Pet1* neuron activity increases the probability of maintaining the active locomotor state.

We next examined whether activation of ROb/Pa *Pet1* neurons influenced locomotor speed. We measured the fraction of time mice spent in defined 10cm/s speed bins and found that ChR mice spent less time engaged in locomotion at slower speeds (<25cm/s) and an increased time at faster speeds (>35cm/s) (Figure 5J). A histogram of time across wheel speed shows a rightward shift during the light periods as compared to no light periods (Figures 5K, 5L, S7D, and S7E), indicating that faster running by ChR mice occurs during light periods. To more precisely determine when light activation influenced speed, we measured average speed during each of the 5 min time bins. We found that average speed increased significantly in ChR animals compared to control animals during the third light period (Figures 5M, S7H, and S7I). We also observed significant increases in the fraction of time spent running >35cm/s and the maximum speed during the third light period compared to the first light period (Figures 5N and 5O). The smaller increases in speed and duration of locomotion during the second light period are maintained during the subsequent no-light period (Figures 5M, 5N, and 5O), suggesting that serotonin continues to influence motor circuits beyond periods of increased neuronal activity. The effects of light activation on locomotion therefore accumulate over time and are strongest during the final light activation period. These observations indicate that descending serotonergic modulation of spinal circuits can influence the speed of locomotion and may be important for regulating behavior over longer time scales.

## Discussion

Neuromodulators are critical to produce flexible and adaptive motor behaviors. While serotonin is a known modulator of spinal circuit dynamics^22^, how the serotonergic system is integrated with motor circuits and how it influences movement has remained unknown. Our findings reveal that spinal cord-projecting serotonergic neurons are interconnected with motor circuits involved in the initiation and execution of locomotion. Serotonergic neurons targeting rhythm-generating spinal circuits receive input from locomotor control areas, display increased neuronal activity during locomotion, and can produce long-lasting potentiation of locomotor behavior. We discuss these findings in the context of neuromodulatory circuit architecture and the function of the serotonergic system in motor control.

Anatomical and molecular genetic neuronal tracing studies have demonstrated the existence of distinct serotonergic populations targeting dorsal and ventral spinal cord^32–38^. Using genetic and viral tracing assays, we provide a more complete picture of the input-output organization of the descending serotonergic system. Our studies show that ventral spinal cord-projecting *Pet1* neurons form a highly branched system that innervates multiple spinal segments, yet in a spatially restricted manner along the dorsoventral domain. The distributed output of descending ROb/Pa *Pet1* neurons likely coordinates modulation of multiple circuits to enable activation of muscles across the body required for behavioral control, including the respiratory and autonomic systems^37,38,58–60^.

We found that ROb/Pa *Pet1* neurons receive input from locomotor control regions including LPGi and CnF. This organization suggests that locomotor command circuits recruit the descending serotonergic system to facilitate changes in excitability required to modulate motor behaviors. Consistent with this model, previous work demonstrated that electrical stimulation of the MLR, including CnF, causes activation of brainstem serotonergic neurons and release of 5-HT in the spinal cord^61,62^. In addition to locomotor-related inputs, we find that ROb/Pa *Pet1* neurons receive input from additional regions that may be essential in mediating context-dependent behaviors. For example, input from ventrolateral PAG, a region known to be activated during defensive freezing responses, may ensure spinal motor circuits are in an appropriate state to produce the required motor response^55,63,64^.

Like other neuromodulators, serotonin acts through volume transmission, but also through direct synaptic contacts^44^. Our transsynaptic tracing studies suggest that spinal MNs receive synaptic input from 5-HT neurons originating from ROb, and to a lesser extent the neighboring caudal 5-HT nuclei. Interestingly, results from our retrograde transsynaptic tracing studies suggest that excitatory V2a INs receive little direct 5-HT synaptic input, as few TPH2+ neurons were rabies-labeled despite robust IN starter cell infection. This suggests possible distinct mechanisms for serotonergic modulation of spinal INs and MNs. Additional layers of specificity are likely to be conferred by differences in 5-HT receptor expression among spinal neuronal classes^65,66^.

Raphe neurons exhibit activity related to levels of tonic motor activity^67^. During REM sleep when there is reduced muscle tone the activity of raphe serotonergic neurons is strongly suppressed, whereas their activity is elevated during waking states. Raphe populations differ however in their activities during specific motor behaviors, such as locomotion. Single-unit recordings in cat showed that the activity of ROb and RPa neurons increases during treadmill locomotion^42^, while activity within DRN neurons remains unchanged^48^. Here, we find that during spontaneous bouts of locomotion, ROb and RPa neurons increase their activity, while DRN neurons decrease activity. While our findings differ from DRN recordings in cat^48^, possibly a result of methodological differences, they are consistent with Ca2+ imagining in mice during open field locomotion, which also show a negative correlation^47^. In addition, our filter analysis of DRN and ROb/Pa data revealed that these relationships between neural activity and locomotion are consistent throughout the spontaneous locomotor behavior.

Anti-correlated activity has also been observed in DRN during locomotion within an open field, yet DRN activity changes in situations where animals perceive a threat, becoming highly correlated to muscle activity^47^. Our findings suggest that similar context-dependent changes in activity occur within dorsal spinal cord projecting RMg neurons. During wheel running we observed that RMg *Pet1* neuron activity was variable with a slight trend towards being anticorrelated with movement. By contrast, during treadmill running RMg *Pet1* neuron activity was strongly correlated with locomotor activity. The forced motorized treadmill assay is likely a more stressful condition for mice compared to unrestrained wheel running. The differences in RMg activity we observed therefore could reflect differences in the internal state of mice in these conditions. RMg modulates incoming sensory information^68–70^, and therefore may act as a context-dependent gating system for sensory feedback. One could imagine a high-threat scenario where high-speed locomotion must be prioritized to promote escape and survival. In this context, it would be distracting or even life-threatening to respond to peripheral nociceptive inputs. In contrast, during non-threatening situations, it would be important to notice and respond to a painful stimulus. In the future it will be important to examine the possible role of RMg in context-dependent gating of sensory feedback and to gauge its influence on motor behavior.

Serotonin modulates the excitability of spinal MNs and INs, generating changes in the temporal dynamics of motor output and magnitude and timing of muscle activation. It has been proposed that serotonin mediates gain control, adjusting the input-output gain of MNs to achieve the desired activation of muscles for specific movements^8,25^. We find that activation of ventral spinal cord-projecting *Pet1* neurons increases the speed and length of running. Activation of serotonergic neurons may function like turning up the gain “knob”, where enhanced release of 5-HT increases the excitability of spinal MNs and INs. This could allow specific motor commands to yield larger or more sustained motor output. Furthermore, gain may differentially influence specific limb muscles for intended motor output, as we have observed variation in 5-HT input across motor pools. Activation of ROb/Pa neurons could also act by sustaining serotonergic modulation, as ROb/Pa neurons have been shown to decrease their firing rates after prolonged locomotion^71^. Conversely, turning down this knob would result in changes to locomotor behavior in the opposite direction, perhaps reducing locomotor duration and speed. Consistent with this model, local delivery of a serotonin receptor antagonist to the lumbar spinal cord results in impaired hindlimb stepping^7^.

Finally, our studies reveal an unappreciated way in which neuromodulatory systems influence locomotion. The descending serotonergic pathway impacts locomotor behavior in a manner that fundamentally differs from the relatively immediate effects that reticulospinal pathways have on locomotor initiation, speed, and termination^51,53,56,57,72^. While the activity of ROb/Pa is tightly synced with locomotor behavior, we find that the neurons do not act as a “go-signal” for locomotion, consistent with what has been observed for the lateral LPGi serotonin neurons^57^. Rather, serotonergic input strengthens and promotes maintenance of ongoing locomotion. We find that activation of ROb/Pa produces increases in locomotor speed and duration, and that these changes accumulate and are sustained over minutes. The influence that the serotonergic pathway has on movement is slower and extends well-beyond the current locomotor bout. These results are reminiscent of work in *C. elegans* showing that 5-HT mediates a long-lasting effect on locomotor speed^5^, suggesting that 5-HT mediates the effects we have observed on locomotion. This enduring influence on behavior may be facilitated by metabotropic 5-HT receptors that result in sustained changes in the excitability of spinal motor circuits. These may be critical for producing sustained locomotor output to meet demands of context-specific motor behaviors.

## Acknowledgements

We thank Kathy Nagel, Michael Long, and Niels Ringstad for valuable discussions and comments on the manuscript. Thank you to Arkarup Banerjee, Dayu Lin, Andrew Miri, Anders Nelson, and Nic Tritsch for helpful discussions and technical advice throughout this work. We thank members of the Lin, Long, Nagel, Schoppik, and Tritsch labs for technical support and advice. Thank you to Pratik Mistry of the Tritsch lab for MATLAB code to perform signal demodulation and baselining for fiber photometry data analysis. We are grateful to Susan Morton for generating the TPH2 antibody, Sebastian Poliak for generating the N2cG-TVA mice, Kim Ritola and Julia Sable for virus preparations, Sarah Pfennig for technical support, and Bryan Chadwick for coding support. Thank you to Nikos Balaskas, Joriene De Nooij, and Andy Murray for technical training and support. Research was supported by the National Institute of Neurological Disorders and Stroke under awards 1K99NS118052 to S.J.F. and 5R35NS116858 to J.S.D., and by the Simons Foundation to S.J.F.

## Author Contributions

S.J.F., J.S.D. and T.M.J. conceived and supervised the project. S.J.F., A.V., A.C., and C.C. performed experiments. S.J.F. performed data analysis. H.G. generated MATLAB code. S.M.D shared mice and provided feedback during collaboration. S.J.F. and J.M.D. wrote manuscript.

## Materials and Methods

### Experimental animals

Male and female mice aged 8-10 weeks at the time of surgical procedures were used for experiments. All mice were maintained on a C57BL/6 genetic background. Prior to surgical procedures, mice were housed communally with littermates. Following surgical procedures and for the duration of behavioral analysis, mice were single housed. Mice were given *ad libitum* access to food and water and maintained on a 12-hour light-dark cycle. All experimental and surgical procedures involving animals were approved by the NYU Grossman School of Medicine Institutional Animals Care and Use Committee (IACUC) and in compliance with the NIH guidelines for care and use of animals.

A listing of all mouse strains using in experiments can be found below in Table 1. To perform monosynaptic retrograde rabies tracing from genetically-defined subpopulations of serotonin neurons (Figure 4), we generated two mouse lines to enable recombinase-dependent expression of the N2c rabies virus glycoprotein (N2cG), and the receptor for the subgroup A avian sarcoma and leukosis virus (TVA). An HA-tagged N2cG and mutated TVA66T^73,74^ were inserted into both the Ai9 (Addgene plasmid #22799) and Ai65 (Addgene plasmid #61577) targeting vector to enable either Cre-dependent or Cre- and Flpe-dependent expression of N2cG and TVA, respectively. Both targeting vectors allow expression of the inserted cassette at the mouse Rosa26 locus. The targeting vectors were used to generate mice at Kallyope and then transferred to NYU Langone animal facility.

**Table 1.** Mouse strains by experiment.

| <u>Experiment</u> | <u>Mouse Strain</u> | <u>Source</u> |
| --- | --- | --- |
| Anterograde labeling of Tac1-Pet1 or Egr2-Pet1 synaptic terminals (Figure 1) | RC::FPSit (Cre- and Flpe-dependent synaptophysin-EGFP)<br>Pet1-Flpe<br>Tac1-Cre<br>Egr2-Cre | S. Dymecki (Harvard)<sup>37,38,43</sup><br>JAX 030206 |
| Monosynaptic rabies virus tracing from spinal motor neurons (Figure 1) | ChAT-Cre | T. Jessell (Columbia)<sup>75</sup><br>Jax 006410 |
| Monosynaptic rabies virus tracing from Chx10<sup>+</sup> interneurons (Figure S1) | Chx10-Cre | T. Jessell (Columbia)<sup>76</sup> |
| Optogenetics and photometry (Figures 2 and 5) | Pet1-Cre | JAX 012712 |
| Monosynaptic rabies tracing from raphe (Figures 4 and S6) | Ai65-N2cG-TVA (Cre- and Flpe-dependent)<br>Ai9-N2cG-TVA (Cre-dependent) | Generated with Sebastian Poliak (Kallyope) |

### Viral injections

Anesthesia was induced using vaporized isoflurane at 3% in oxygen (2 L/min). Anesthesia was maintained throughout the duration of procedures at 1.5-2.5% in oxygen (1 L/min) with mice held in stereotaxic frame (Kopf Instruments, Model 940) atop a feedback-controlled heating pad set to 37 °C. Viruses were injected using a Nanoject II (Drummond Scientific Company) and pulled glass capillaries.

To perform rabies tracing from spinal MNs, AAV9-Syn-DIO-TVA66T-tdTomato-N2cG (Julie Sable, Jessell lab, Columbia) was injected into the lateral ventricle by intracerebroventricular (ICV) injection at P0-1. The ICV injection allows virus to spread within the cerebrospinal fluid and enables infection of spinal MNs^46^. At eight weeks of age, mice were injected with EnvA-N2c(ΔG)-TdT (Janelia, NeuroTools Viral Vector Core) into spinal segments C4 to T1. To perform rabies tracing from Ch10+ INs, Chx10-Cre mice were injected with AAV1-flex-TVA-H2B-HA-N2cG (Julie Sable, Jessell lab, Columbia) into spinal segments C5 to C8, and three weeks later EnvA-N2c(ΔG)-TdT (Julie Sable, Jessell lab, Columbia) was injected into segments C5 to C8. Mice were perfused 1-2 weeks post rabies injection. To retrogradely label serotonergic neurons from the spinal cord (Figure S3), AAV2retro-CAG-FLEX-tdTomato (Addgene 28306-AAVrg) was injected into spinal segments C5 to C8 of Pet1-cre mice (8 weeks). Mice were perfused 2 weeks post-injection. For rabies tracing in *N2cG-TVA* mice, EnvA-N2c(ΔG)-TdT (Janelia, NeuroTools Viral Vector Core) was injected using ROb coordinates described below and mice were perfused 7-8 days post-injection.

Coordinates for ROb injections were 6.85mm posterior and 0mm mediolateral of bregma with injection at depth of 5.4 to 5.7mm below bregma. Coordinates for RMg injections were 5.3mm posterior and 0mm mediolateral of bregma with injection at depth of 5.7mm. Coordinates for DRN injections were 4.7mm posterior and 0mm mediolateral of bregma with injection at depth of 3mm. Cell-type specific expression of Gcamp6s and channelrhodopsin was achieved using the following Cre-dependent viruses: AAV5-FLEX-GcAMP6s (Addgene #100842-AAV5) for ROb/Pa and RMg, AAV1-FLEX-GcAMP6s (Addgene #100842-AAV1) for DRN, and AAV2-EF1a-DIO-ChR2(H134R)-EYFP (UNC Vector Core). Approximately 300 nL of virus was injected per animal. Following the delivery of viral vector, a 26-gauge guide cannula (P1 Technologies) was positioned overtop of the injection site, descended to a depth of .2mm above desired fiber position and removed. A fiberoptic cannula (400µm core diameter, Doric Lenses) was then lower into the brain and cemented in place (C&B Metabond, Parkell). Lastly, a headplate was cemented to the skull to facilitate handling of mice while attaching and removing the fiber optic cable.

To retrogradely label spinal motor neurons targeting the gluteus muscle, ChAT-Cre mice (n=3) were injected with AAV2r-flex-tdTomato (Addgene #28306-AAVrg) at age P5-6. Pups were anesthetized using isoflurane and 1-2uL of virus was injected into the gluteus muscle. Mice were perfused 3 weeks post injection and used for immunostaining and analysis of 5-HT fibers in spinal cord sections.

### Histology

Mice were euthanized via isoflurane overdose delivered by the open drop method and transcardially perfused with 10mL of cold 1X phosphate buffered saline (PBS) followed by 10mL of ice cold 4% paraformaldehyde (PFA). Brain and/or spinal cords were removed and post-fixed overnight in 4% PFA at 4°C. Tissue was sectioned free floating in cold 1X PBS using a vibratome (Leica VT1200S). Brain tissue was sectioned at 70 μm thickness and spinal cord tissue was sectioned at 80 μm thickness.

For immunohistochemistry, sections were permeabilized in 0.3% TritonX-100 in 1X PBS for 15 minutes at room temperature. Following permeabilization, sections were incubated free floating in primary antibodies (Table 2) diluted in a blocking solution of 1% bovine serum albumin (BSA) and 0.3% TritonX-100 in 1X PBS for 72 hours at 4°C. Secondary antibodies (Table 3) diluted in blocking solution were applied overnight at 4°C. Sections were mounted onto superfrost plus microscope slides (Fisherbrand) with Fluoromount-G (SouthernBiotech).

**Table 2:**
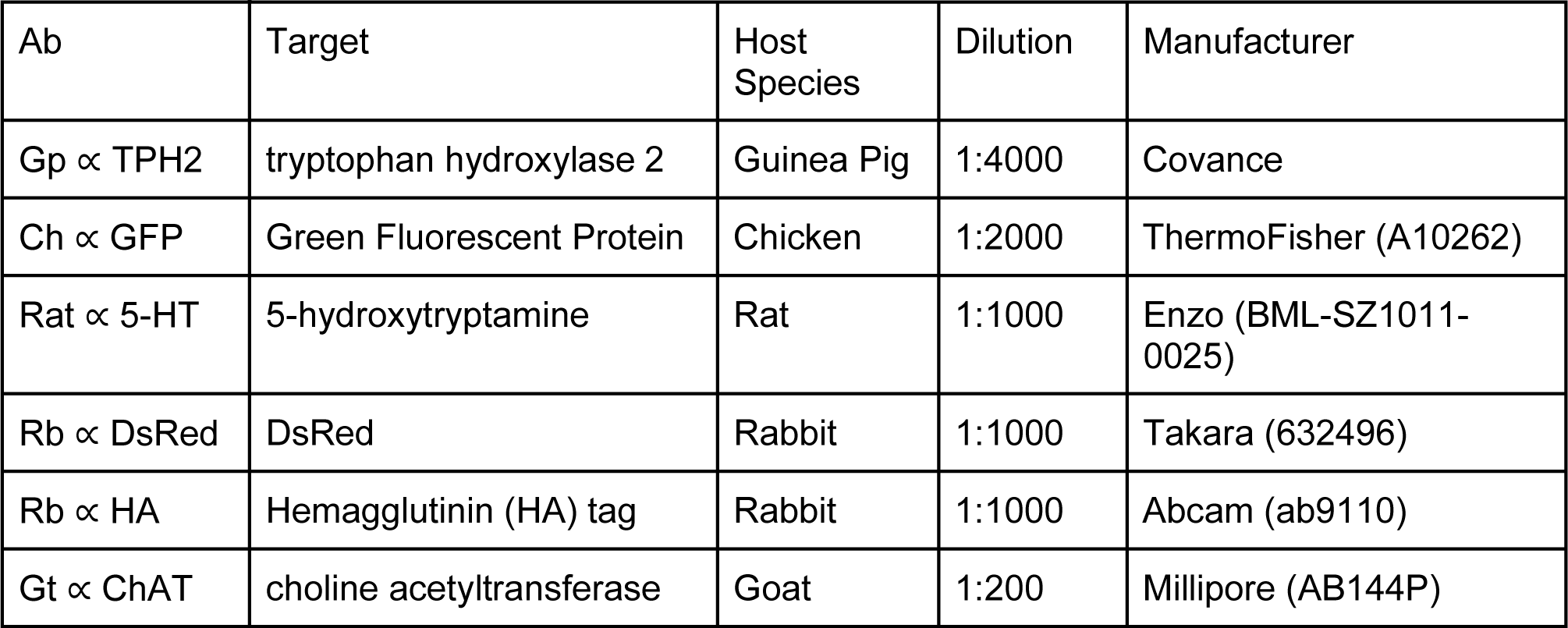
Primary Antibodies.

**Table 3:** Secondary Antibodies.

| Ab | Conjugated fluorophore | Dilution | Manufacturer & Cat # |
| --- | --- | --- | --- |
| Donkey $\alpha$ Guinea Pig - Cy3 | Cyanine 3 | 1:1000 | Jackson (706-165-148) |
| Donkey $\alpha$ Guinea Pig - 647 | Alexa Fluor 647 | 1:1000 | Jackson (706-605-148) |
| Donkey $\alpha$ Chicken - 488 | Alexa Fluor 488 | 1:1000 | Jackson (703-545-155) |
| Donkey $\alpha$ Rat - Cy3 | Cyanine 3 | 1:1000 | Jackson (712-165-150) |
| Donkey $\alpha$ Rat - Cy5 | Cyanine 5 | 1:1000 | Jackson (712-175-150) |
| Donkey $\alpha$ Rabbit - Cy3 | Cyanine 3 | 1:1000 | Jackson (711-165-152) |
| Donkey $\alpha$ Rabbit - 488 | Alexa Fluor 488 | 1:1000 | Jackson (711-545-152) |
| Donkey $\alpha$ Goat - 647 | Alexa Fluor 647 | 1:1000 | Jackson (705-605-147) |

TPH2 antibody was generated by Covance using a TPH2 peptide (RRGLSLDSAVPEDHQLC, Atlantic Peptides) coupled to KLH (Thermofisher 77605). Reagents were designed and prepared by Susan Brenner-Morton at Columbia University.

### Image analysis

Imaris (Bitplane) software was used to analyze the position of synaptophysinGFP puncta in spinal cord sections and MATLAB was used to generate the position plots and density plots for each 5-HT neuron population, as previously described in Bikoff et al.^77^. Single images at each spinal level were used to generate the position and density plots for *Egr2-Pet1* and *Tac1-Pet1* puncta, although these patterns were reproducible across at least 3 animals examined. This approach was also used to analyze 5-HT puncta and gluteal MN density in ChAT-Cre mice. For this experiment, position data was combined from 3 lumbar spinal cord images from each of 3 injected animals to generate density plots. Additionally, this approach was used to generate the position plots of RV+ and TPH2+ cells following rabies tracing from spinal MNs and Chx10+ INs.

### Fluorescent *In Situ* hybridization

*In situ* hybridization was performed on brain tissue following the injection of CVS-N2c(ΔG)-tdT rabies virus to identify retrogradely labeled cells expressing the vesicular glutamate transporter 2 (Vglut2). Hybridization chain reaction (HCR) probes for Vglut2 were generated from the sense sequence of Vglut2 cDNA (NCBI) using the HCR 3.0 probe maker^78,79^. Following tissue collection and post-fixation described above, brains were placed in a 30% sucrose solution overnight at 4 °C, then embedded in Tissue-Tek O.C.T. compound. Tissue was cryosectioned at 18µm thickness, collected onto superfrost plus microscope slides, and stored at -80 °C. Fluorescent in situ hybridization was performed using the Hybridization Chain Reaction RNA fluorescent in situ hybridization (HCR RNA-FISH, Molecular Instruments) protocol, described in D’Elia et al.^80^.

### Locomotor assays and acclimation

Wheel running experiments were performed using a low-profile running wheel (Fast Trac K3251, Bio-Serve) attached to rotary encoder (A2 optical shaft encoder, US Digital). Encoder output was collected using a USB interface board (RHD2000, #C3100, Intan Technologies) and recorded using RHD2000 Interface software (Intan Technologies). Treadmill experiments were performed using a custom-built motorized rodent treadmill (Model 802, University of Cologne electronics lab).

Mice were acclimated to the running wheel and treadmill for 4 days prior to the first behavioral data collection trials. On the first day of behavioral acclimation, mice were allowed explore the wheel and treadmill environments for 10 minutes each. On the subsequent 3 days, mice were acclimated with the fiber optic cable attached. Each of these 3 days, mice were then allowed to run freely on the wheel for 10-30 minutes. On the second day of behavioral acclimation, mice ran on the treadmill for 1 minute at 20 cm/sec and 1 minute at 25 cm/sec with a 1-minute rest period in between. The treadmill speed was ramped up slowly by hand to acclimate the animal to the moving belt. On the third and fourth days of behavioral acclimation, mice on the treadmill performed alternating 10 second intervals of rests and runs: 3X at 20 cm/sec, 3X at 25 cm/sec, 3X at 30 cm/sec, and 3X at 35 cm/sec.

### Fiber photometry

A rig for collecting fiber photometry data was constructed using LEDs (M470F3 & M405FP1, Thorlabs), Fluorescent Mini Cube (FMC Gen1, Doric), and photoreceiver (Model 2151, Newport). A 385 Hz sinusoidal 470 LED light (30µW, bandpass filtered 460-490nm) was delivered by a 400µm optic cable to excite Gcamp6s in the brain. A 315 Hz sinusoidal 405 LED light (30µW, bandpass filtered 400-410nm) was also delivered to the brain as a control for motion artifact. The isosbestic point for Gcamp6s is near 405nm, where emitted light is independent of calcium binding. The emitted light from the brain was bandpass filtered (500-550nm) and collected by photodetector and recorded using the Intan USB interface board with Intan software described above with locomotor assays.

Recordings were made during 30min wheel sessions and treadmill runs over 14-21 days and collected at 4kHz sampling rate. The signals were bandpass filtered (470nm passing band: 385Hz ± 10, 405 passing band: 315Hz ± 10 and demodulated using a phase sensitive detection method comparing the modulated signal with a recorded reference signal of same frequency. Demodulation extracts the envelope of the 385Hz signal reflecting the intensity of Gcamp6s signal. A baseline was calculated from demodulated signal using an interpolated linear fit of values in 10^th^ percentile and moving window size of 30s. The photometry signal was baseline adjusted: ΔF/F= (demodulated signal-baseline)/baseline.

The averaged ΔF/F at the start and stop of run bouts was generated using runs of at least 3 seconds in length, defined by periods where wheel speed was greater than 10cm/sec. For this analysis, each run was baselined individually. Run starts were normalized to the average fluorescence 0.5s to 1s prior to the run start and run stops to the fluorescence 0.5s to 1s post run stop. To analyze the relationship between neural activity and speed, the averaged ΔF/F and wheel speed were plotted for the early phase of each run and throughout each run. The early phase was defined as seconds 2 and 3 of each run. The full run was defined as the entire run without the first and last seconds of each run. The beginning and end of each run were not used in the analysis to remove acceleration and deceleration phases of the behavior, so to isolate the period of sustained running when comparing activity with speed. Animals that displayed no signal and where histology confirmed insufficient Gcamp expression or misplacement of fiber position were excluded from our analysis.

### Filter calculation and analysis

Filters were calculated as described in Nagel and Doupe, 2006^49^. Filters were extracted using concatenated ΔF/F and wheel speed data for each individual animal. An average filter and SEM were generated for each brain area (n=3 for each area: ROb, RMg, and DRN). The filter was used to predict locomotor speed from ΔF/F for each brain area and then compared to the actual wheel speed. A correlation coefficient was calculated for each animal to determine the strength of the relationship between the predicted speed and actual speed. An average correlation coefficient and SEM were calculated for each brain area.

### Optogenetic activation and analysis

For optical stimulation of ChR2, 470nm LED light (M470F3, Thorlabs) was delivered for 5s pulses at 25Hz with sinusoidal temporal profile. Light intensity was measured using an optical power meter (PM100D, Thorlabs) at the tip of the fiber canula to be 13-15 mW. During each of the 5-minute light periods, mice received 15 5-second light pulses.

Mice were attached to optic fiber and allowed to run freely on wheel for 30min during the light stimulation protocol with 5-minute alternating no light and light periods. Data was collected from each ChR and control mice once a day with 1-2 rest days in between for 3 weeks. For data to be included in analysis, the following criteria were required: 1) at least 5 minutes total of speeds >5cm/sec and 2) reached speeds greater than 10cm/sec during all 3 no light and light periods. These criteria were used to ensure comparable behavior when animals ran to a similar extent throughout the 30 minutes and remove days where mice stopped running for long periods of time. For wheel speed analysis, wheel rotary encoder output was converted to speed using the wheel circumference and maximum encoder output value and downsampled to 1kHz. Run bouts were defined as periods of wheel speed greater than 10cm/sec for at least 1 second.

### Electromyographic recordings and analysis

To observe the relationship between 5-HT neuron activity and muscle activity during locomotor behavior, electromyographic (EMG) electrodes were implanted into limb muscle to record muscle activity. EMG electrodes were fabricated and implanted as previously described^81^. Briefly, EMG electrodes were made using insulated steel wire (793200, A-M Systems) with 1mm exposed regions 7.5cm away from a 12-pin miniature connector (11P3828, Newark). The wire ends were inserted and crimped inside of a 27-gauge needle to use when inserting electrodes into the muscle. EMG electrodes were implanted into mice that had previously been injected with Gcamp6s virus and had headplates attached. An incision was made above the neck, hip and tibialis anterior (TA) muscle in right hindlimb. The needle end of the wire was guided beneath the skin from the neck to the hip and then to TA. The needle was used to insert the wire through the muscle and a knot made at the end to hold the electrode in place. The connecter was cemented to the rear edge of the headplate. EMG signals were transmitted from the headplate connecter by an Omnetics connector and amplified using an amplifier chip (RHD2216, Intan Technologies). Amplified EMG signals were recorded using the Intan USB interface board and Intan software. EMG recordings were downsampled to 1kHz, high-pass filtered at 40Hz, and rectified, as described in Miri et al^81^.

---

**Figure S1.**
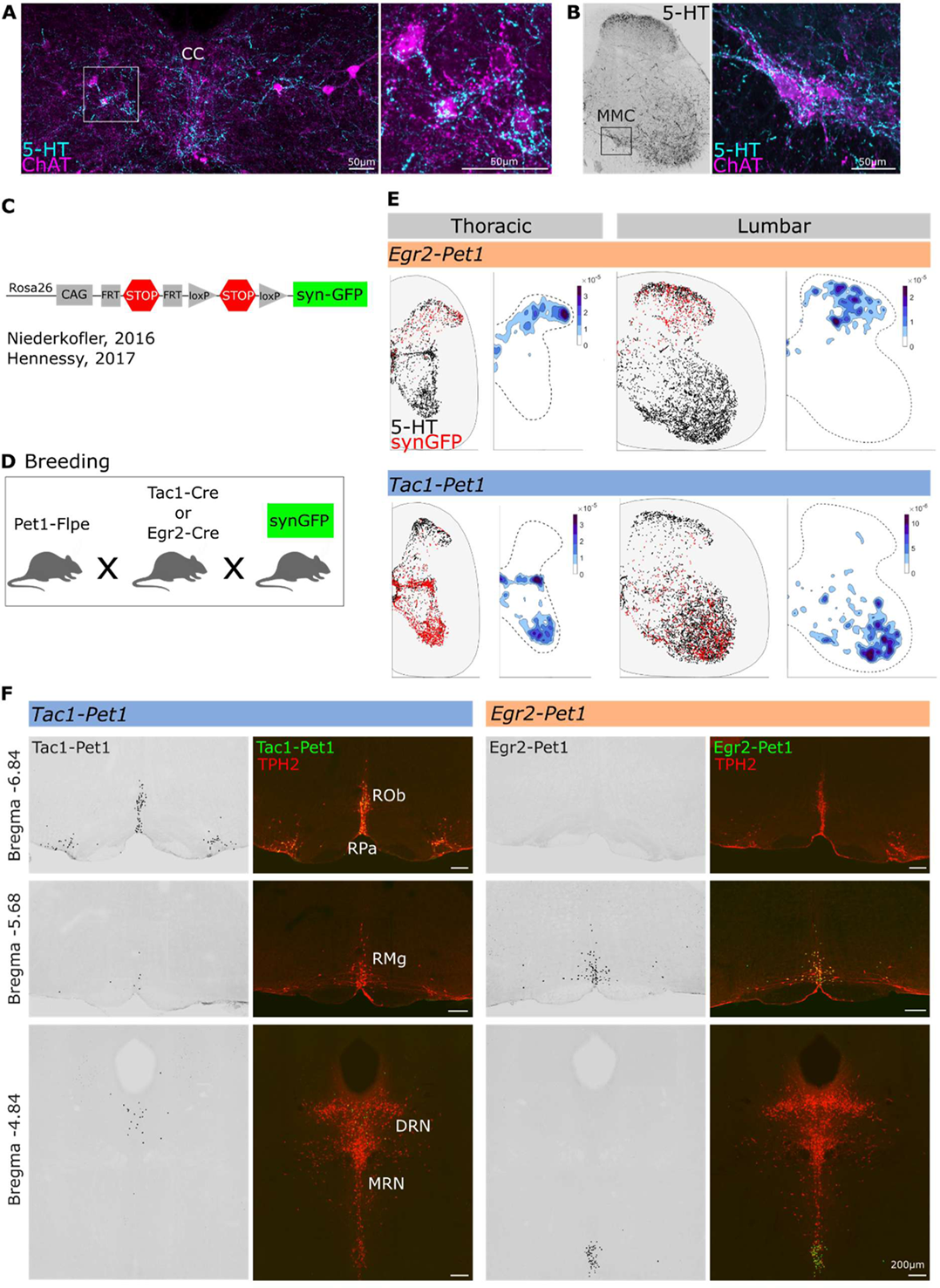
Genetic labeling of 5-HT sub-populations. **(A)** 5-HT immunostaining of cervical spinal cord showing 5-HT puncta at cholinergic interneurons beside central canal (CC). Higher magnification image on right of area within white box. Scale bars 50µm. **(B)** 5-HT immunostaining at MMC (black box) in lumbar spinal cord with higher magnification of motor neurons on right. Scale bar 50µm. **(C).** Intersectional synaptophysin-GFP allele ^38,43^. **(D)** Breeding scheme to generate *Egr2-Pet1-synGFP* or *Tac1-Pet1-synGFP* mice. **(E)** Distribution of synaptophysinGFP (red) and 5-HT immunostaining (black) puncta in thoracic and lumbar spinal segments for *Egr2-Pet1* and *Tac1-Pet1* neuron populations. Relative density of synGFP puncta (blue). Cervical segments in Figure 1. **(F)** Distribution of *Tac1-Pet1* and *Egr2-Pet1* neurons in brainstem and midbrain with TPH2 immunostaining. Scale bars 200µm. Raphe pallidus, RPa; raphe obscurus, ROb; raphe magnus, RMg; dorsal raphe nucleus, DRN; median raphe nucleus, MRN.

**Figure S2.**
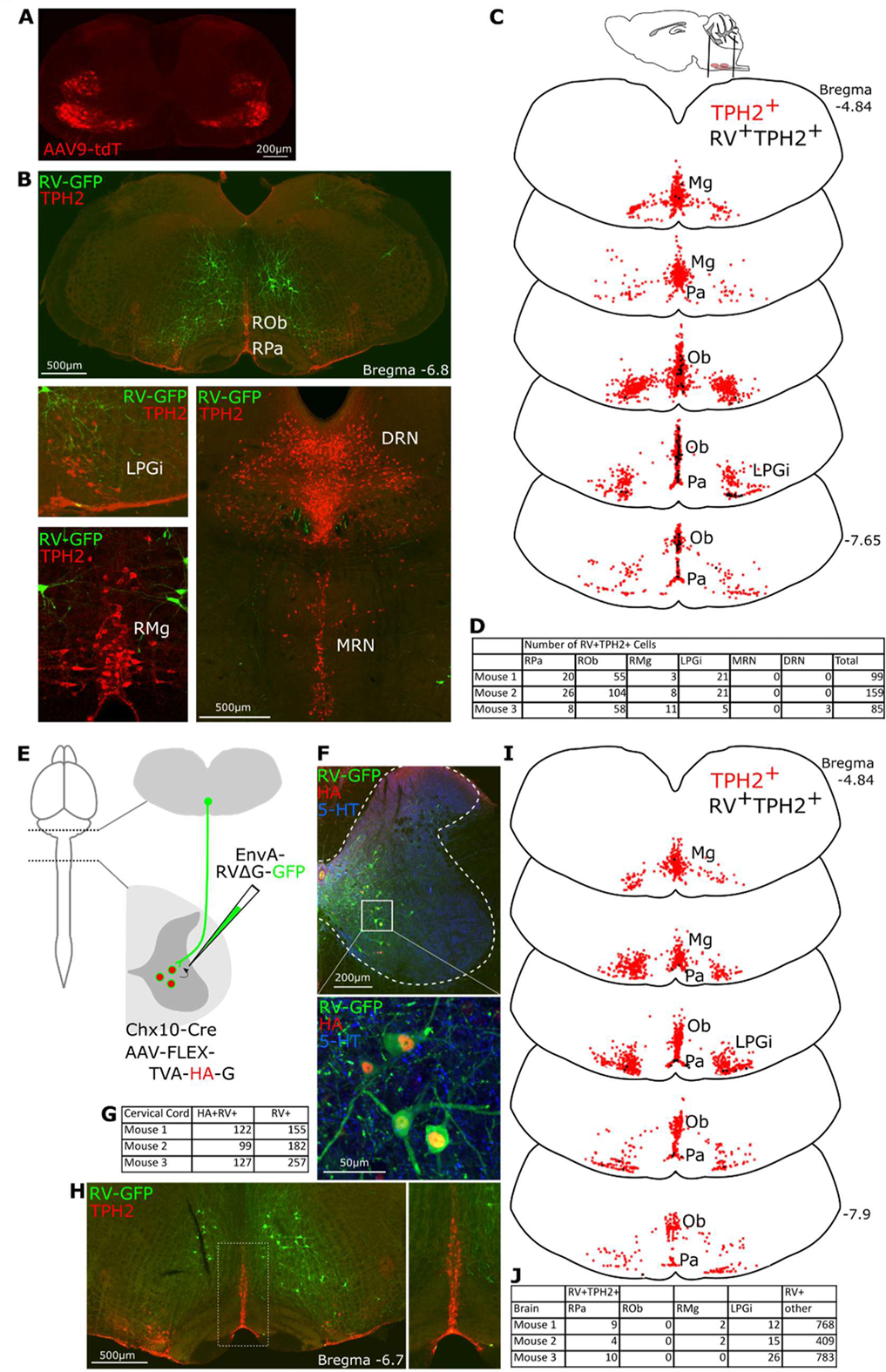
Monosynaptic rabies tracing from spinal MNs and Chx10^+^ Ins. **(A)** MN-restricted expression of tdTomato (tdT) in spinal cord (C8) following intracerebraoventricular injection (ICV) of Cre-dependent AAV9 virus into ChAT-Cre mouse. **(B)** Rabies virus (RV) labeling in brainstem and midbrain with TPH2 immunostaining showing serotonergic nuclei. Scale bars 500µm. **(C)** Summary of RV^+^TPH2^+^ neurons in single *ChAT-Cre* animal with total TPH2^+^ cells. **(D)** Cell counts of second-order RV+ TPH2+ neurons in raphe nuclei by animal. **(E)** Experimental strategy for tracing monosynaptic inputs to Chx10^+^ INs. Injection of *Chx10-cre* mouse cervical spinal cord (C5-C8) with AAV-FLEX-TVA-HA-G followed by EnvA-RVΔG-GFP. **(F)** Infection of Chx10^+^ neurons in cervical spinal cord (C4). Scale bar 200µm. Zoom-in below showing RV-infected HA^+^ neurons and 5-HT puncta. Scale bar 50µm. **(G)** Cell counts of primary infected HA+RV+ neurons and second order RV+ (HA-) neurons in cervical spinal cord by animal. **(H)** RV labeling within brainstem reticular formation. Scale bar 500µm. Image on right shows magnified ROb/Pa region **(I)** Summary of RV^+^TPH2^+^ neurons in single *Chx10-cre* animal with total TPH2^+^ cells. **(J)** Cell counts of second-order RV+TPH2+ neurons in caudal raphe and LPGi regions and second-order RV+TPH2-neurons throughout the rest of the brainstem. (Raphe pallidus, RPa; raphe obscurus, ROb; lateral paragigantocellularis, LPGi; raphe magnus, RMg; dorsal raphe nucleus, DRN; median raphe nucleus, MRN.)

**Figure S3.**
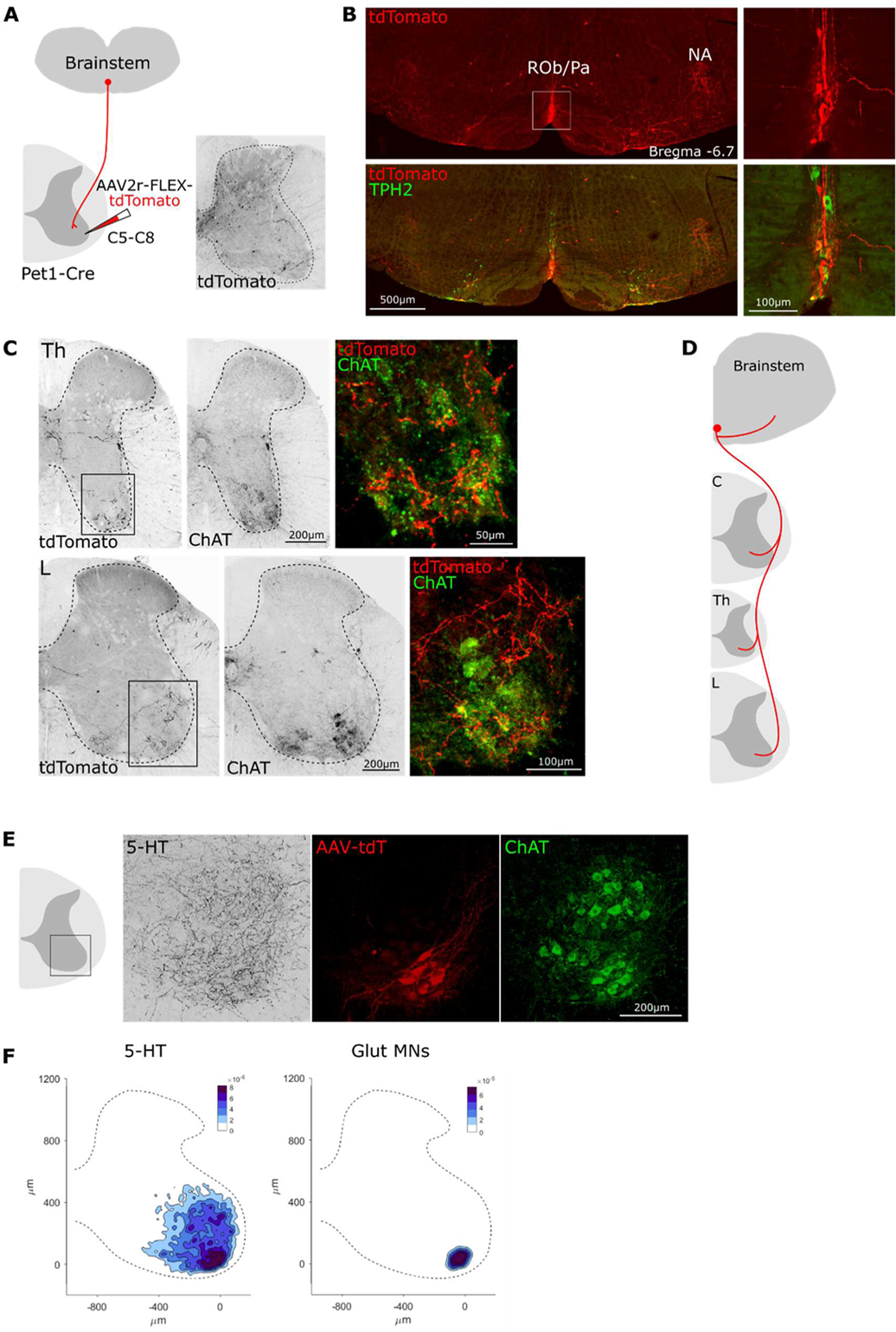
Retrograde labeling of spinal-projecting *Pet1* neurons. **(A)** Experimental procedure. Injection of AAV2r-FLEX-tdTomato into C5-C8 of *Pet1-Cre* mice. Expression of tdTomato at injected cervical level. **(B)** Expression of tdTomato in brainstem with TPH2 expression. tdTomato^+^ fibers at nucleus ambiguous (NA). Scale bar 500µm. Zoom-in of expression within white box. Scale bar 100µm. **(C)** tdTomato expression within fibers of thoracic (Th) and lumbar (L) spinal cord with ChAT immunostaining. Right most images are zoom-in of black box showing tdTomato^+^ fibers around MNs. **(D)** Schematic depicting highly collateralized ROb/Pa *Pet1* neurons targeting multiple spinal levels and caudal medulla. **(E)** Expression of 5-HT in lumbar ventral horn with retrogradely infected gluteal MNs (AAV-tdT). Scale bar 200µm. **(F)** Relative density plots for ventral horn 5-HT puncta and gluteal MNs (n=3 animals).

**Figure S4.**
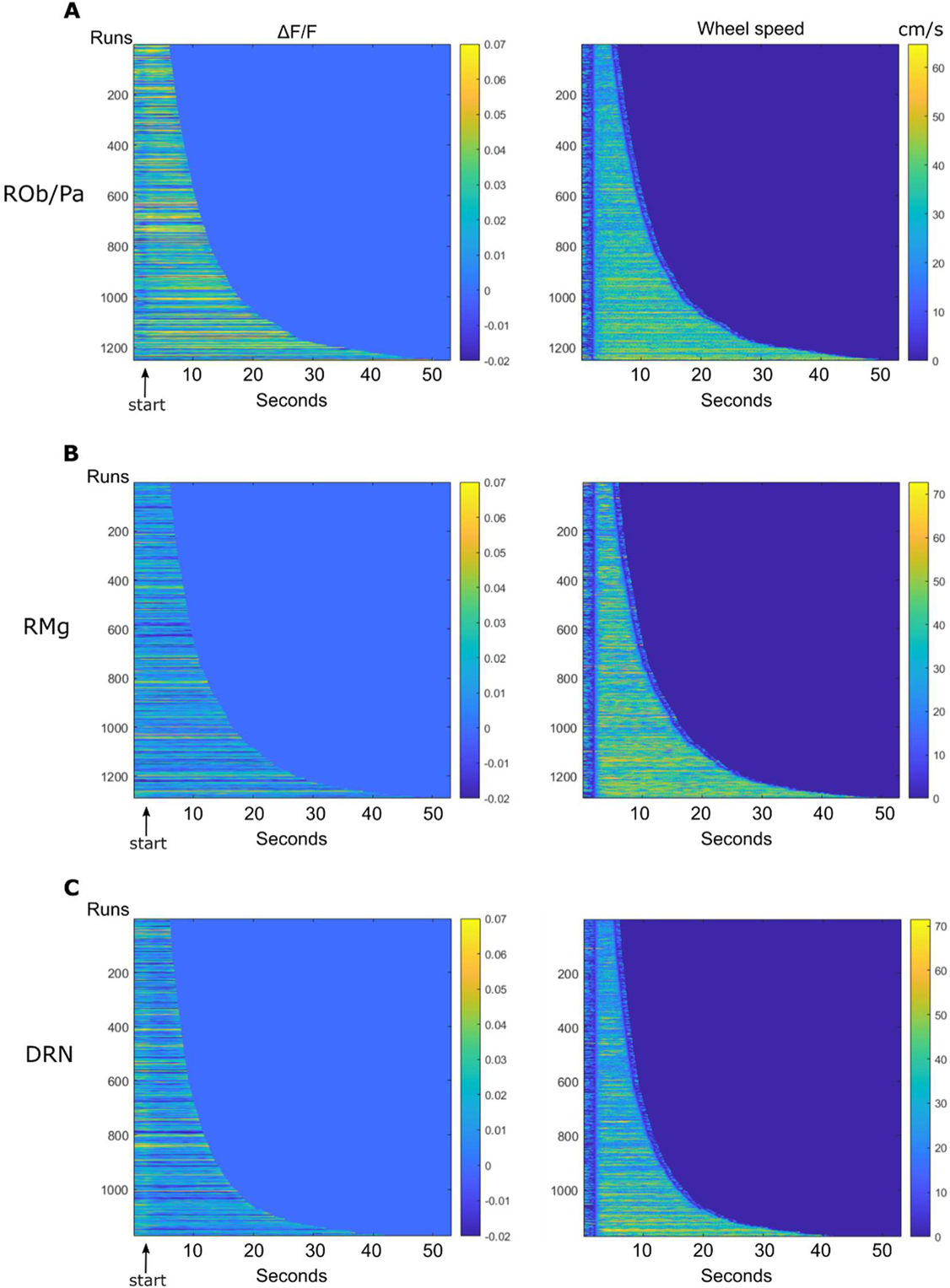
Neural activity during individual runs on wheel (A-C) Heatmap showing ΔF/F (left) for all individual runs during wheel assay. *Pet1* neuron activity within ROb/Pa, RMg, or DRN. Each horizontal line is a single run. Runs ordered by length from shortest to longest. Traces include 2 seconds prior to run start. Heatmap showing corresponding wheel speed (right) for all runs. **(A)** ROb/Pa (1252 runs, 4 animals). **(B)** RMg (1290 runs, 4 animals). **(C)** DRN (1172 runs, 3 animals).

**Figure S5.**
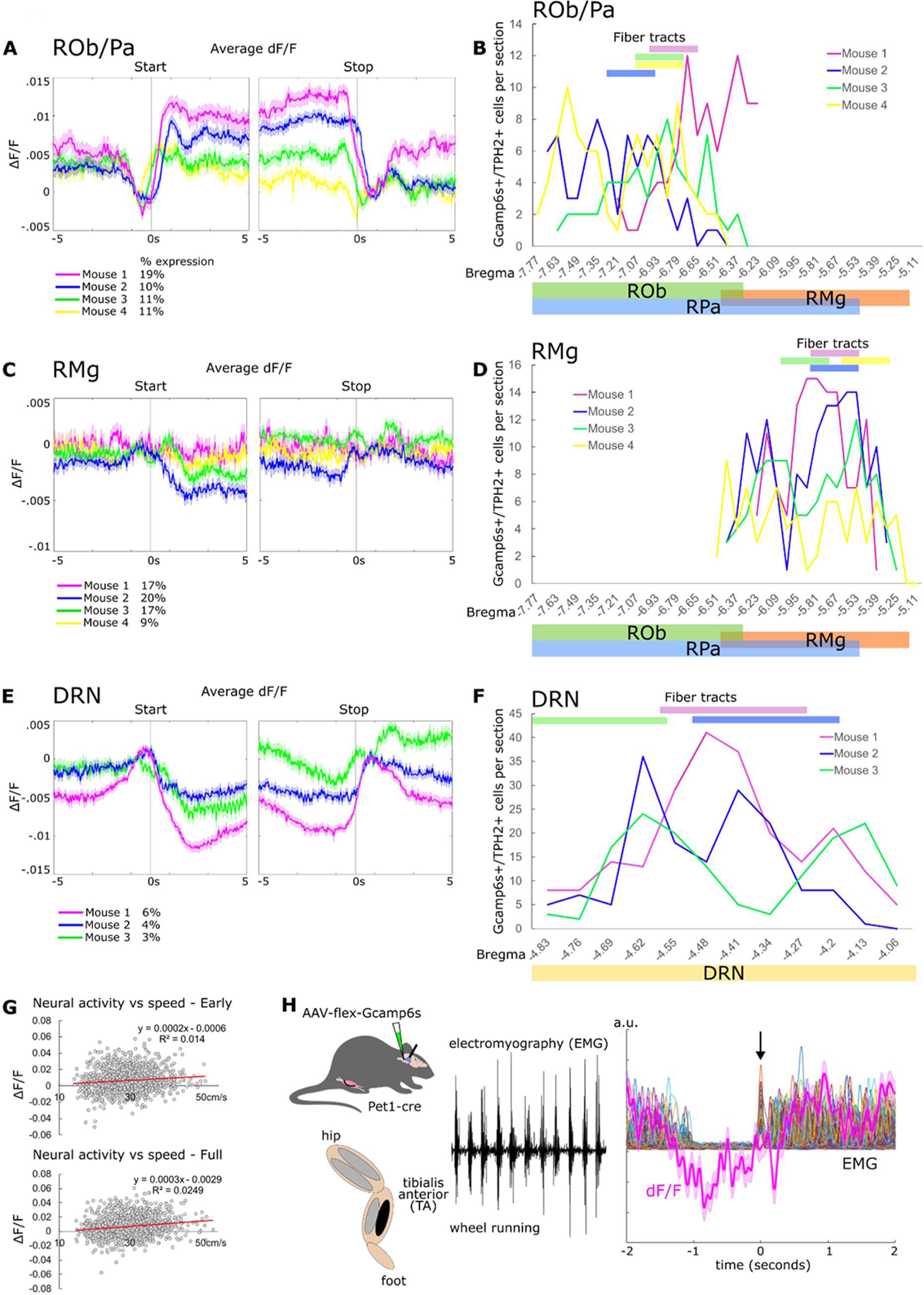
Additional photometry data and histology quantification (A,C,E) Averaged 470nm ΔF/F by animal at run start and stop. Percentage of Gcamp6s-infected ROb/Pa TPH2+ cells is listed below by animal. (A) ROb/Pa. (C) RMg. (E) DRN. **(B,D,F)** Post-hoc quantification of histology for animals used in fiber photometry experiments. Gcamp6s^+^TPH2^+^ cell counts and fiber position for each animal. (B) ROb/Pa. (D) RMg. (F) DRN. **(G)** Plot of averaged ΔF/F and wheel speed during 2sec early in the run (top) and during the full run (bottom). **(H)** EMG recordings from tibialis anterior (TA) with Gcamp imaging during wheel running. Raw EMG trace from TA during locomotion (black). Average ΔF/F (pink) from ROb/Pa overlaying rectified EMG traces aligned to first peak of muscle activity at start of run bout (multicolor). Arrow shows alignment of first muscle burst.

**Figure S6.**
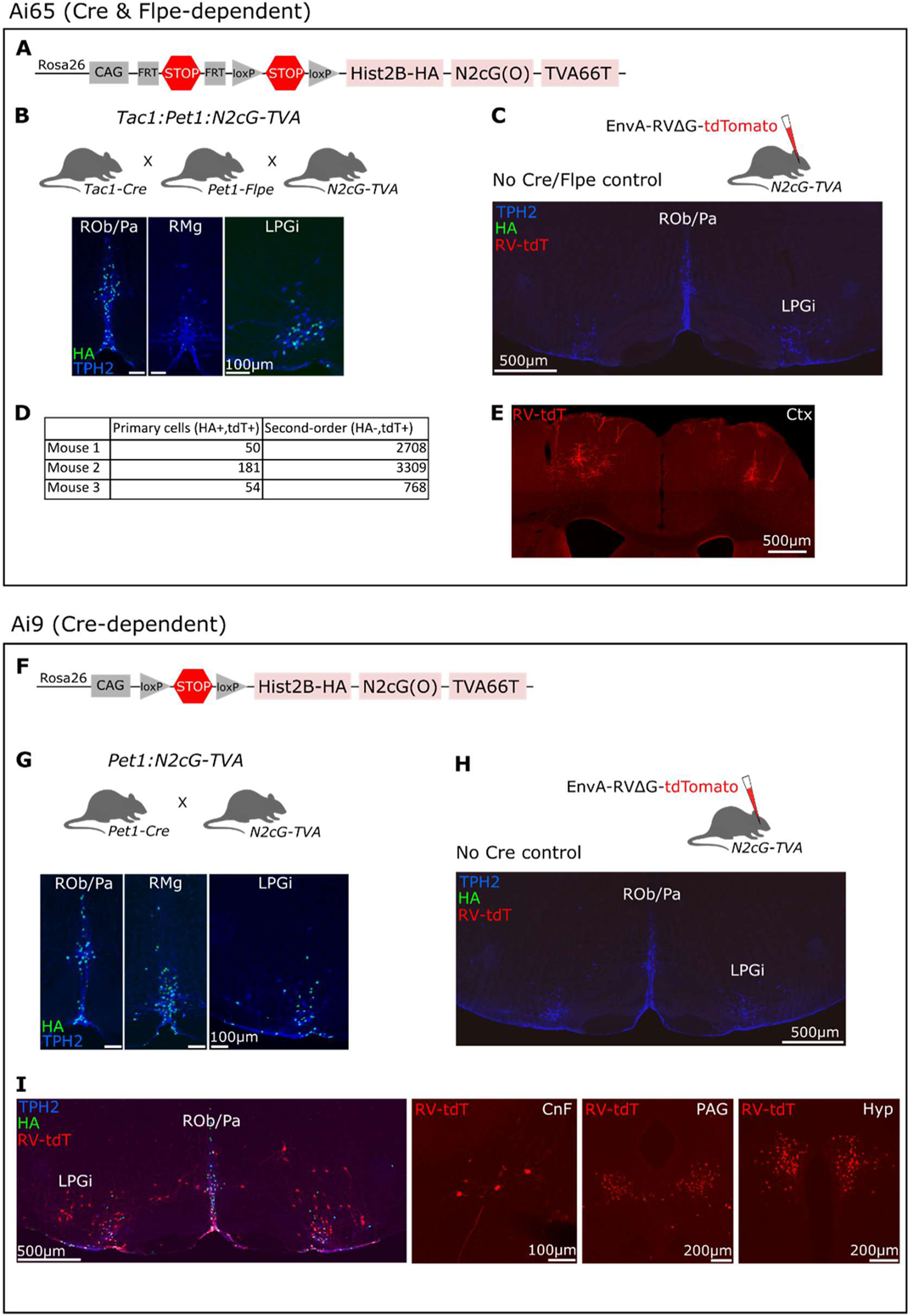
Retrograde rabies tracing from raphe neurons targeting ventral spinal cord. **(A)** Intersectional *N2cG-TVA* allele. **(B)** Generation of *Tac1:Pet1:N2cG-TVA* mice. Expression of HA-tagged N2cG within ROb/Pa and LPGi neurons. **(C)** HA and TPH2 immunostaining in *N2cG-TVA* mouse (no Cre/Flpe) injected with EnvA-RVΔG-tdTomato. **(D)** Cell counts of primary infected cells (HA+tdT+) and second-order (HA-tdT+) cells by animal from *Tac1:Pet1:N2cG-TVA* mice injected with EnvA-RVΔG-tdTomato. **(E)** Second order rabies-labeled neurons in sensorimotor cortex (Ctx) **(F)** Cre-dependent *N2cG-TVA* allele. **(G)** Generation of *Pet1:N2cG-TVA* mice. Expression of HA-tagged N2cG within ROb/Pa, RMg, LPGi neurons. **(H)** HA and TPH2 immunostaining in *N2cG-TVA* mouse (no Cre) injected with EnvA-RVΔG-tdTomato. **(I)** Rabies infection in *Pet1:N2cG-TVA* caudal brainstem with labeled presynaptic cells in LPGi, cuneiform (CnF), periaqueductal grey (PAG), and hypothalamus (Hyp).

**Figure S7.**
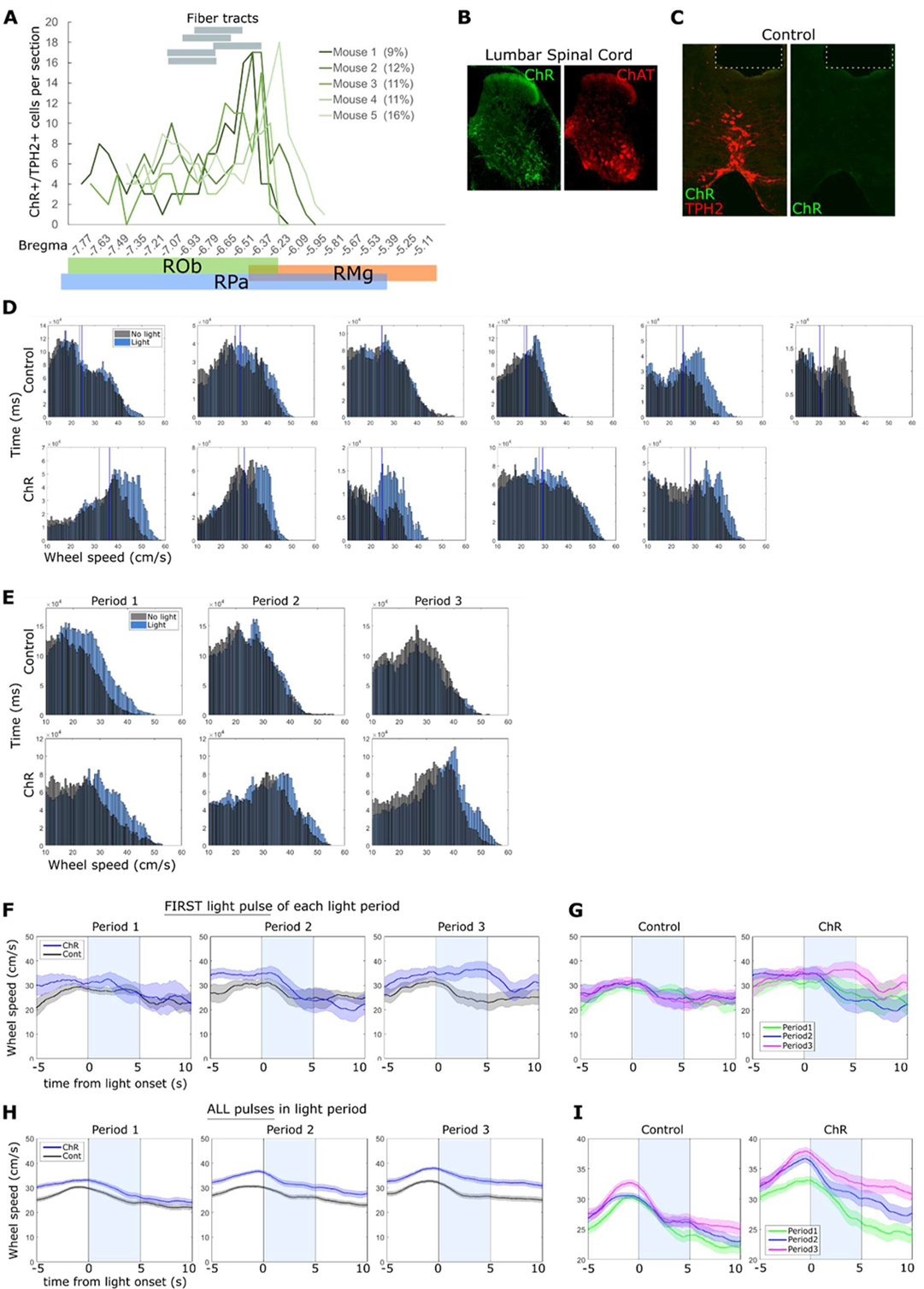
Optogenetic activation of ROb/Pa *Pet1* neurons during locomotor behavior. **(A)** ChR-infected cells and fiber position for each ChR animal. Percentage of ChR-infected ROb/Pa TPH2+ cells is listed beside each animal. **(B)** ChR expression in lumbar spinal cord with ChAT immunostaining. **(C)** Representative control animal with no ChR expression. White dotted line denotes position of fiber. **(D)** Distribution of time (ms) across wheel speeds during No-light (grey) or Light (blue) periods (1-3) shown for each individual Control (top row) or ChR (bottom row) animal. **(E)** Distribution of time (ms) across wheel speeds during No-light (grey) or Light (blue) periods (1-3). Combined data from 5-6 animals. **(F)** Averaged wheel speed when animal was running at time of light onset. Averaged wheel speed during first light pulse of each period when wheel speed was greater than 20cm/s at light onset. **(G)** Data in F displayed by control or ChR animals with light periods overlayed. **(H)** Averaged wheel speed across all light pulses of each period when wheel speed was greater than 20cm/s at light onset. **(I)** Data in H displayed by control or ChR animals with overlayed light periods.

## References

1. Katz, P.S. (1995). Intrinsic and extrinsic neuromodulation of motor circuits. Curr Opin Neurobiol 5, 799–808. 10.1016/0959-4388(95)80109-x.

2. Marder, E. (2012). Neuromodulation of neuronal circuits: back to the future. Neuron 76, 1–11. 10.1016/j.neuron.2012.09.010.

3. Marder, E., and Bucher, D. (2001). Central pattern generators and the control of rhythmic movements. Curr Biol 11, R986–996. 10.1016/s0960-9822(01)00581-4.

4. Harris-Warrick, R.M. (2011). Neuromodulation and flexibility in Central Pattern Generator networks. Curr Opin Neurobiol 21, 685–692. 10.1016/j.conb.2011.05.011.

5. Huang, Y.C., Luo, J., Huang, W., Baker, C.M., Gomes, M.A., Meng, B., Byrne, A.B., and Flavell, S.W. (2023). A single neuron in C. elegans orchestrates multiple motor outputs through parallel modes of transmission. Curr Biol 33, 4430–4445 e4436. 10.1016/j.cub.2023.08.088.

6. Howard, C.E., Chen, C.-L., Tabachnik, T., Hormigo, R., Ramdya, P., and Mann, R.S. (2019). Serotonergic Modulation of Walking in Drosophila. Current Biology 29, 4218--4230.e4218. 10.1016/j.cub.2019.10.042, pmid = 31786064, pmcid = PMC6935052.

7. Cabaj, A.M., Majczynski, H., Couto, E., Gardiner, P.F., Stecina, K., Slawinska, U., and Jordan, L.M. (2017). Serotonin controls initiation of locomotion and afferent modulation of coordination via 5-HT7 receptors in adult rats. J Physiol 595, 301--320. 10.1113/jp272271, pmid = 27393215.

8. Wei, K., Glaser, J.I., Deng, L., Thompson, C.K., Stevenson, I.H., Wang, Q., Hornby, T.G., Heckman, C.J., and Kording, K.P. (2014). Serotonin Affects Movement Gain Control in the Spinal Cord. The Journal of Neuroscience 34, 12690--12700. 10.1523/jneurosci.1855-14.2014, pmid = 25232107, pmcid = PMC4166156.

9. Lillvis, J.L., and Katz, P.S. (2013). Parallel evolution of serotonergic neuromodulation underlies independent evolution of rhythmic motor behavior. J Neurosci 33, 2709–2717. 10.1523/JNEUROSCI.4196-12.2013.

10. Dunbar, M.J., Tran, M.A., and Whelan, P.J. (2010). Endogenous extracellular serotonin modulates the spinal locomotor network of the neonatal mouse. J Physiol 588, 139–156. 10.1113/jphysiol.2009.177378.

11. Perrier, J.F., and Delgado-Lezama, R. (2005). Synaptic release of serotonin induced by stimulation of the raphe nucleus promotes plateau potentials in spinal motoneurons of the adult turtle. J Neurosci 25, 7993--7999. 10.1523/jneurosci.1957-05.2005, pmid = 16135756.

12. Clemens, S., and Katz, P.S. (2001). Identified serotonergic neurons in the Tritonia swim CPG activate both ionotropic and metabotropic receptors. J Neurophysiol 85, 476–479. 10.1152/jn.2001.85.1.476.

13. Zhang, W., and Grillner, S. (2000). The spinal 5-HT system contributes to the generation of fictive locomotion in lamprey. Brain Res 879, 188–192. 10.1016/s0006-8993(00)02747-5.

14. Kiehn, O., and Kjaerulff, O. (1996). Spatiotemporal characteristics of 5-HT and dopamine-induced rhythmic hindlimb activity in the in vitro neonatal rat. J Neurophysiol 75, 1472–1482. 10.1152/jn.1996.75.4.1472.

15. Barbeau, H., and Rossignol, S. (1991). Initiation and modulation of the locomotor pattern in the adult chronic spinal cat by noradrenergic, serotonergic and dopaminergic drugs. Brain Res 546, 250--260. 10.1016/0006-8993(91)91489-n, pmid = 2070262.

16. Katz, P.S., and Harris-Warrick, R.M. (1990). Neuromodulation of the crab pyloric central pattern generator by serotonergic/cholinergic proprioceptive afferents. J Neurosci 10, 1495–1512. 10.1523/JNEUROSCI.10-05-01495.1990.

17. Wallen, P., Christenson, J., Brodin, L., Hill, R., Lansner, A., and Grillner, S. (1989). Mechanisms underlying the serotonergic modulation of the spinal circuitry for locomotion in lamprey. Prog Brain Res 80, 321–327; discussion 315-329. 10.1016/s0079-6123(08)62227-x.

18. Hounsgaard, J., and Kiehn, O. (1989). Serotonin-induced bistability of turtle motoneurones caused by a nifedipine-sensitive calcium plateau potential. J Physiol 414, 265–282. 10.1113/jphysiol.1989.sp017687.

19. Beltz, B., Eisen, J.S., Flamm, R., Harris-Warrick, R.M., Hooper, S.L., and Marder, E. (1984). Serotonergic innervation and modulation of the stomatogastric ganglion of three decapod crustaceans (Panulirus interruptus, Homarus americanus and Cancer irroratus). J Exp Biol 109, 35–54. 10.1242/jeb.109.1.35.

20. Livingstone, M.S., Harris-Warrick, R.M., and Kravitz, E.A. (1980). Serotonin and octopamine produce opposite postures in lobsters. Science 208, 76–79. 10.1126/science.208.4439.76.

21. Flaive, A., Fougere, M., van der Zouwen, C.I., and Ryczko, D. (2020). Serotonergic Modulation of Locomotor Activity From Basal Vertebrates to Mammals. Front Neural Circuits 14, 590299. 10.3389/fncir.2020.590299.

22. Perrier, J.F., and Cotel, F. (2015). Serotonergic modulation of spinal motor control. Curr Opin Neurobiol 33, 1--7. 10.1016/j.conb.2014.12.008, pmid = 25553359.

23. Cazalets, J.R., Sqalli-Houssaini, Y., and Clarac, F. (1992). Activation of the central pattern generators for locomotion by serotonin and excitatory amino acids in neonatal rat. J Physiol 455, 187–204. 10.1113/jphysiol.1992.sp019296.

24. Barbeau, H., and Rossignol, S. (1990). The effects of serotonergic drugs on the locomotor pattern and on cutaneous reflexes of the adult chronic spinal cat. Brain Research 514, 55--67. 10.1016/0006-8993(90)90435-e, pmid = 2357531.

25. Johnson, M.D., and Heckman, C.J. (2014). Gain control mechanisms in spinal motoneurons. Front Neural Circuits 8, 81. 10.3389/fncir.2014.00081, pmid = 25120435, pmcid = PMC4114207.

26. Hounsgaard, J., and Kiehn, O. (1985). Ca++ dependent bistability induced by serotonin in spinal motoneurons. Exp Brain Res 57, 422--425. 10.1007/bf00236551, pmid = 2578974.

27. Husch, A., Dietz, S.B., Hong, D.N., and Harris-Warrick, R.M. (2015). Adult spinal V2a interneurons show increased excitability and serotonin-dependent bistability. J Neurophysiol 113, 1124–1134. 10.1152/jn.00741.2014.

28. Perrier, J.F., and Hounsgaard, J. (2003). 5-HT2 receptors promote plateau potentials in turtle spinal motoneurons by facilitating an L-type calcium current. J Neurophysiol 89, 954--959. 10.1152/jn.00753.2002, pmid = 12574471.

29. Steinbusch, H.W.M. (1981). Distribution of serotonin-immunoreactivity in the central nervous system of the rat-cell bodies and terminals. Neuroscience 6, 557--618. 10.1016/0306-4522(81)90146-9, pmid = 7017455.

30. Dahlstroem, A., and Fuxe, K. (1964). Evidence for the Existence of Monoamine-Containing Neurons in the Central Nervous System. I. Demonstration of Monoamines in the Cell Bodies of Brain Stem Neurons. Acta Physiol Scand Suppl, SUPPL 232:231–255.

31. Hornung, J.P. (2003). The human raphe nuclei and the serotonergic system. J Chem Neuroanat 26, 331–343. 10.1016/j.jchemneu.2003.10.002.

32. Basbaum, A.I., Clanton, C.H., and Fields, H.L. (1978). Three bulbospinal pathways from the rostral medulla of the cat: An autoradiographic study of pain modulating systems. Journal of Comparative Neurology 178 209--224. 10.1002/cne.901780203, pmid = 627624.

33. Holstege, G., and Kuypers, H.G.J.M. (1982). The Anatomy of Brain Stem Pathways to the Spinal Cord in Cat. A Labeled Amino Acid Tracing Study. Progress in Brain Research 57, 145--175. 10.1016/s0079-6123(08)64128-x, pmid = 7156396.

34. Martin, R.F., Jordan, L.M., and Willis, W.D. (1978). Differential projections of cat medullary raphe neurons demonstrated by retrograde labelling following spinal cord lesions. Journal of Comparative Neurology 182, 77--88. 10.1002/cne.901820106, pmid = 701490.

35. Loewy, A.D. (1981). Raphe pallidus and raphe obscurus projections to the intermediolateral cell column in the rat. Brain Res 222, 129–133. 10.1016/0006-8993(81)90946-x.

36. Holstege, J.C., and Kuypers, H.G.J.M. (1987). Brainstem projections to spinal motoneurons: An update. Neuroscience 23, 809--821. 10.1016/0306-4522(87)90160-6, pmid = 2893995.

37. Brust, R., Corcoran, A., Richerson, G., Nattie, E., and Dymecki, S. (2014). Functional and developmental identification of a molecular subtype of brain serotonergic neuron specialized to regulate breathing dynamics. Cell Rep 9, 2152--2165. 10.1016/j.celrep.2014.11.027, pmid = 25497093, pmcid = PMC4351711.

38. Hennessy, M., Corcoran, A., Brust, R., Chang, Y., Nattie, E., and Dymecki, S. (2017). Activity of Tachykinin1-Expressing Pet1 Raphe Neurons Modulates the Respiratory Chemoreflex. J Neurosci 37, 1807--1819. 10.1523/jneurosci.2316-16.2016, pmid = 28073937.

39. Okaty, B., Freret, M., Rood, B., Brust, R., Hennessy, M., deBairos, D., Kim, J., Cook, M., and Dymecki, S. (2015). Multi-Scale Molecular Deconstruction of the Serotonin Neuron System. Neuron 88, 774--791. 10.1016/j.neuron.2015.10.007, pmid = 26549332, pmcid = PMC4809055.

40. Okaty, B., Commons, K., and Dymecki, S. (2019). Embracing diversity in the 5-HT neuronal system. Nat Rev Neurosci 20, 397--424. 10.1038/s41583-019-0151-3, pmid = 30948838.

41. Okaty, B.W., Sturrock, N., Escobedo Lozoya, Y., Chang, Y., Senft, R.A., Lyon, K.A., Alekseyenko, O.V., and Dymecki, S.M. (2020). A single-cell transcriptomic and anatomic atlas of mouse dorsal raphe Pet1 neurons. Elife 9. 10.7554/eLife.55523.

42. Veasey, S.C., Fornal, C.A., Metzler, C.W., and Jacobs, B.L. (1995). Response of serotonergic caudal raphe neurons in relation to specific motor activities in freely moving cats. The Journal of Neuroscience 15, 5346--5359. 10.1523/jneurosci.15-07-05346.1995, pmid = 7623157, pmcid = PMC6577863.

43. Niederkofler, V., Asher, T.E., Okaty, B.W., Rood, B.D., Narayan, A., Hwa, L.S., Beck, S.G., Miczek, K.A., and Dymecki, S.M. (2016). Identification of Serotonergic Neuronal Modules that Affect Aggressive Behavior. Cell Rep 17, 1934–1949. 10.1016/j.celrep.2016.10.063.

44. Alvarez, F.J., Pearson, J.C., Harrington, D., Dewey, D., Torbeck, L., and Fyffe, R.E. (1998). Distribution of 5-hydroxytryptamine-immunoreactive boutons on alpha-motoneurons in the lumbar spinal cord of adult cats. J Comp Neurol 393, 69–83.

45. Reardon, T.R., Murray, A.J., Turi, G.F., Wirblich, C., Croce, K.R., Schnell, M.J., Jessell, T.M., and Losonczy, A. (2016). Rabies Virus CVS-N2c(DeltaG) Strain Enhances Retrograde Synaptic Transfer and Neuronal Viability. Neuron 89, 711–724. 10.1016/j.neuron.2016.01.004.

46. Simon, C.M., Dai, Y., Van Alstyne, M., Koutsioumpa, C., Pagiazitis, J.G., Chalif, J.I., Wang, X., Rabinowitz, J.E., Henderson, C.E., Pellizzoni, L., and Mentis, G.Z. (2017). Converging Mechanisms of p53 Activation Drive Motor Neuron Degeneration in Spinal Muscular Atrophy. Cell Rep 21, 3767–3780. 10.1016/j.celrep.2017.12.003.

47. Seo, C., Guru, A., Jin, M., Ito, B., Sleezer, B.J., Ho, Y.Y., Wang, E., Boada, C., Krupa, N.A., Kullakanda, D.S., et al. (2019). Intense threat switches dorsal raphe serotonin neurons to a paradoxical operational mode. Science 363, 538–542. 10.1126/science.aau8722.

48. Veasey, S.C., Fornal, C.A., Metzler, C.W., and Jacobs, B.L. (1997). Single-unit responses of serotonergic dorsal raphe neurons to specific motor challenges in freely moving cats. Neuroscience 79, 161–169. 10.1016/s0306-4522(96)00673-2.

49. Nagel, K.I., and Doupe, A.J. (2006). Temporal processing and adaptation in the songbird auditory forebrain. Neuron 51, 845–859. 10.1016/j.neuron.2006.08.030.

50. Dayan, P.A., L.F. (2001). Theoretical Neuroscience (The MIT Press).

51. Capelli, P., Pivetta, C., Esposito, M.S., and Arber, S. (2017). Locomotor speed control circuits in the caudal brainstem. Nature 551, 373--377. 10.1038/nature24064, pmid = 29059682.

52. Caggiano, V., Leiras, R., Goñi-Erro, H., Masini, D., Bellardita, C., Bouvier, J., Caldeira, V., Fisone, G., and Kiehn, O. (2018). Midbrain circuits that set locomotor speed and gait selection. Nature 553, 455--460. 10.1038/nature25448, pmid = 29342142, pmcid = PMC5937258.

53. Ferreira-Pinto, M.J., Ruder, L., Capelli, P., and Arber, S. (2018). Connecting Circuits for Supraspinal Control of Locomotion. Neuron 100, 361--374. 10.1016/j.neuron.2018.09.015, pmid = 30359602.

54. Berg, E.M., Mrowka, L., Bertuzzi, M., Madrid, D., Picton, L.D., and El Manira, A. (2023). Brainstem circuits encoding start, speed, and duration of swimming in adult zebrafish. Neuron 111, 372–386 e374. 10.1016/j.neuron.2022.10.034.

55. Tovote, P., Esposito, M.S., Botta, P., Chaudun, F., Fadok, J.P., Markovic, M., Wolff, S.B., Ramakrishnan, C., Fenno, L., Deisseroth, K., et al. (2016). Midbrain circuits for defensive behaviour. Nature 534, 206–212. 10.1038/nature17996.

56. Leiras, R., Cregg, J.M., and Kiehn, O. (2022). Brainstem Circuits for Locomotion. Annual Review of Neuroscience 45, 1--23. 10.1146/annurev-neuro-082321-025137, pmid = 34985919.

57. Hsu, L.-J., Bertho, M., and Kiehn, O. (2023). Deconstructing the modular organization and real-time dynamics of mammalian spinal locomotor networks. Nature Communications 14, 873. 10.1038/s41467-023-36587-w, pmid = 36797254, pmcid = PMC9935527.

58. DePuy, S.D., Kanbar, R., Coates, M.B., Stornetta, R.L., and Guyenet, P.G. (2011). Control of Breathing by Raphe Obscurus Serotonergic Neurons in Mice. The Journal of Neuroscience 31, 1981--1990. 10.1523/jneurosci.4639-10.2011, pmid = 21307236, pmcid = PMC3071248.

59. Bacon, S.J., Zagon, A., and Smith, A.D. (1990). Electron microscopic evidence of a monosynaptic pathway between cells in the caudal raphe nuclei and sympathetic preganglionic neurons in the rat spinal cord. Exp Brain Res 79, 589–602. 10.1007/BF00229327.

60. Lindsay, A.D., and Feldman, J.L. (1993). Modulation of respiratory activity of neonatal rat phrenic motoneurones by serotonin. J Physiol 461, 213–233. 10.1113/jphysiol.1993.sp019510.

61. Noga, B.R., Turkson, R.P., Xie, S., Taberner, A., Pinzon, A., and Hentall, I.D. (2017). Monoamine Release in the Cat Lumbar Spinal Cord during Fictive Locomotion Evoked by the Mesencephalic Locomotor Region. Front Neural Circuits 11, 59. 10.3389/fncir.2017.00059.

62. Opris, I., Dai, X., Johnson, D.M.G., Sanchez, F.J., Villamil, L.M., Xie, S., Lee-Hauser, C.R., Chang, S., Jordan, L.M., and Noga, B.R. (2019). Activation of Brainstem Neurons During Mesencephalic Locomotor Region-Evoked Locomotion in the Cat. Front Syst Neurosci 13, 69. 10.3389/fnsys.2019.00069.

63. Carrive, P. (1993). The periaqueductal gray and defensive behavior: functional representation and neuronal organization. Behav Brain Res 58, 27–47. 10.1016/0166-4328(93)90088-8.

64. Watson, T.C., Cerminara, N.L., Lumb, B.M., and Apps, R. (2016). Neural Correlates of Fear in the Periaqueductal Gray. J Neurosci 36, 12707–12719. 10.1523/JNEUROSCI.1100-16.2016.

65. Fonseca, M.I., Ni, Y.G., Dunning, D.D., and Miledi, R. (2001). Distribution of serotonin 2A, 2C and 3 receptor mRNA in spinal cord and medulla oblongata. Brain Res Mol Brain Res 89, 11--19. 10.1016/s0169-328x(01)00049-3, pmid = 11311971.

66. Koyama, Y., Kondo, M., and Shimada, S. (2017). Building a 5-HT3A Receptor Expression Map in the Mouse Brain. Sci Rep 7, 42884. 10.1038/srep42884, pmid = 28276429.

67. L, J.B., and A, F.C. (1993). 5-HT and motor control: a hypothesis. Trends Neurosci 16, 346--352. 10.1016/0166-2236(93)90090-9, pmid = 7694403.

68. Basbaum, A.I., Clanton, C.H., and Fields, H.L. (1976). Opiate and stimulus-produced analgesia: functional anatomy of a medullospinal pathway. Proc Natl Acad Sci U S A 73, 4685–4688. 10.1073/pnas.73.12.4685.

69. Cai, Y.Q., Wang, W., Hou, Y.Y., and Pan, Z.Z. (2014). Optogenetic activation of brainstem serotonergic neurons induces persistent pain sensitization. Mol Pain 10, 70. 10.1186/1744-8069-10-70.

70. Ganley, R.P., de Sousa, M.M., Werder, K., Ozturk, T., Mendes, R., Ranucci, M., Wildner, H., and Zeilhofer, H.U. (2023). Targeted anatomical and functional identification of antinociceptive and pronociceptive serotonergic neurons that project to the spinal dorsal horn. Elife 12. 10.7554/eLife.78689.

71. Fornal, C.A., Martin-Cora, F.J., and Jacobs, B.L. (2006). “Fatigue” of medullary but not mesencephalic raphe serotonergic neurons during locomotion in cats. Brain Res 1072, 55–61. 10.1016/j.brainres.2005.12.007.

72. Bouvier, J., Caggiano, V., Leiras, R., Caldeira, V., Bellardita, C., Balueva, K., Fuchs, A., and Kiehn, O. (2015). Descending Command Neurons in the Brainstem that Halt Locomotion. Cell 163, 1191--1203. 10.1016/j.cell.2015.10.074, pmid = 26590422.

73. Callaway, E.M., and Luo, L. (2015). Monosynaptic Circuit Tracing with Glycoprotein-Deleted Rabies Viruses. The Journal of neuroscience : the official journal of the Society for Neuroscience 35, 8979--8985. 10.1523/jneurosci.0409-15.2015, pmid = 26085623, pmcid = PMC4469731.

74. Miyamichi, K., Shlomai-Fuchs, Y., Shu, M., Weissbourd, B.C., Luo, L., and Mizrahi, A. (2013). Dissecting local circuits: parvalbumin interneurons underlie broad feedback control of olfactory bulb output. Neuron 80, 1232–1245. 10.1016/j.neuron.2013.08.027.

75. Balaskas, N., Abbott, L.F., Jessell, T.M., and Ng, D. (2019). Positional Strategies for Connection Specificity and Synaptic Organization in Spinal Sensory-Motor Circuits. Neuron 102, 1143–1156 e1144. 10.1016/j.neuron.2019.04.008.

76. Azim, E., Jiang, J., Alstermark, B., and Jessell, T.M. (2014). Skilled reaching relies on a V2a propriospinal internal copy circuit. Nature 508, 357–363. 10.1038/nature13021.

77. Bikoff, J.B., Gabitto, M.I., Rivard, A.F., Drobac, E., Machado, T.A., Miri, A., Brenner-Morton, S., Famojure, E., Diaz, C., Alvarez, F.J., et al. (2016). Spinal Inhibitory Interneuron Diversity Delineates Variant Motor Microcircuits. Cell 165, 207–219. 10.1016/j.cell.2016.01.027.

78. Choi, H.M.T., Schwarzkopf, M., Fornace, M.E., Acharya, A., Artavanis, G., Stegmaier, J., Cunha, A., and Pierce, N.A. (2018). Third-generation in situ hybridization chain reaction: multiplexed, quantitative, sensitive, versatile, robust. Development 145. 10.1242/dev.165753.

79. Kuehn, E., Clausen, D.S., Null, R.W., Metzger, B.M., Willis, A.D., and Ozpolat, B.D. (2022). Segment number threshold determines juvenile onset of germline cluster expansion in Platynereis dumerilii. J Exp Zool B Mol Dev Evol 338, 225–240. 10.1002/jez.b.23100.

80. D’Elia, K.P., Hameedy, H., Goldblatt, D., Frazel, P., Kriese, M., Zhu, Y., Hamling, K.R., Kawakami, K., Liddelow, S.A., Schoppik, D., and Dasen, J.S. (2023). Determinants of motor neuron functional subtypes important for locomotor speed. Cell Rep 42, 113049. 10.1016/j.celrep.2023.113049.

81. Miri, A., Warriner, C.L., Seely, J.S., Elsayed, G.F., Cunningham, J.P., Churchland, M.M., and Jessell, T.M. (2017). Behaviorally Selective Engagement of Short-Latency Effector Pathways by Motor Cortex. Neuron 95, 683–696 e611. 10.1016/j.neuron.2017.06.042.

